# A molecular network for functional versatility of *HECATE* transcription factors

**DOI:** 10.1101/253815

**Authors:** Christophe Gaillochet, Suraj Jamge, Froukje van der Wal, Gerco Angenent, Richard Immink, Jan U. Lohmann

## Abstract

During the plant life cycle, diverse signalling inputs are continuously integrated and engage specific genetic programs depending on the cellular or developmental context. Consistent with an important role in this process, HECATE (HEC) bHLH transcription factors display diverse functions, from photomorphogenesis to the control of shoot meristem dynamics and gynoecium patterning. However, the molecular mechanisms underlying their functional versatility and the deployment of specific HEC sub-programs still remain elusive.

To address this issue, we systematically identified proteins with the capacity to interact with HEC1, the best characterized member of the family, and integrated this information with our data set of direct HEC1 target genes. The resulting core genetic modules were consistent with specific developmental functions of HEC1, including its described activities in light signalling, gynoecium development and auxin homeostasis. Importantly, we found that in addition, *HEC* genes play a role in the modulation of flowering time and uncovered that their role in gynoecium development may involve the direct transcriptional regulation of *NGATHA1 (NGA1)* and *NGA2* genes. NGA factors were previously shown to contribute to fruit development, but our data now show that they also modulate stem cell homeostasis in the SAM.

Taken together, our results suggest a molecular network underlying the functional versatility of HEC transcription factors. Our analyses have not only allowed us to identify relevant target genes controlling shoot stem cell activity and a so far undescribed biological function of HEC1, but also provide a rich resource for the mechanistic elucidation of further context dependent HEC activities.

**Significance statement:** Although many transcription factors display diverse regulatory functions during plant development, our understanding of the underlying mechanisms remains poor. Here, by reconstructing the regulatory modules orchestrated by the bHLH transcription factor HECATE1 (HEC1), we defined its regulatory signatures and delineated a regulatory network that provides a molecular basis for its functional versatility. In addition, we uncovered a function for *HEC* genes in modulating flowering time and further identified downstream signalling components balancing shoot stem cell activity.

## Introduction

During the continuous elaboration of the plant body, morphogenetic processes take place as gene regulatory networks are deployed in space and time. In particular, phytohormones and transcription factors are orchestrated and trigger specific developmental programs depending on the cellular context (reviewed in (Weijers & Wagner 2016; Schaller et al. 2015)). In line with this concept, the bHLH transcription factors HECATE1 (HEC1), HEC2 and HEC3 control multiple developmental processes throughout the *Arabidopsis thaliana* life cycle in a partially redundant fashion (Zhu et al. 2016; Gremski et al. 2007; Schuster, Gaillochet & Lohmann 2015; Gaillochet et al. 2017).

After germination, seedlings sense light availability and can trigger the establishment of two distinct developmental programs: photomorphogenesis in light, or skotomorphogenesis in dark (reviewed in (Xu et al. 2015)). Two signalling components are crucial during this process: phytochromes that are the photoreceptors perceiving the Red to Far Red light ratio (R/FR), and the PHYTOCHROME INTERACTING FACTOR (PIF) bHLH transcription factors that transcriptionally repress the initiation of photomorphogenesis (reviewed in (Duek & Fankhauser 2005)). Upon light perception at seedling emergence, phytochromes are activated through a conformational change and translocate to the nucleus to repress PIF activities (Leivar & Quail 2011). HEC factors modulate this signalling pathway by forming a protein complex with PIF1 and PIF3 that inhibits their binding to DNA, preventing them from exerting their transcriptional function and consequently positively regulating photomorphogenesis (Zhu et al. 2016).

The integration of light signals is also crucial to trigger the transition from the vegetative to the reproductive phase of development, mainly by sensing day-length (reviewed in (Bäurle & Dean 2006)). Flowering transition is orchestrated by the florigen FLOWERING LOCUS T (FT), which is directly activated by CONSTANS (CO) in phloem companion cells of the leaf (Wigge et al. 2005). Interestingly, the photoreceptors PHYTOCHROME A (PHYA) and PHYB antagonistically control the stability of CO at different time of the day and thus directly influence this developmental transition (Valverde et al. 2004).

Later during development, HEC factors regulate cellular behavior at the shoot apical meristem (SAM) (Schuster et al. 2014; Gaillochet et al. 2017). The shoot meristem is subdivided in different functional domains and harbors stem cells that continuously generate aboveground tissues. The organizing centre (OC) acts as a stem cell niche and maintains stem cell identity in the overlying central zone (CZ) (Mayer et al. 1998; Schoof et al. 2000; Brand et al. 2000). Mechanistically, *WUSCHEL* (WUS) RNA is expressed exclusively in the OC whereas WUS protein moves through plasmodesmata towards the CZ to instruct stem cell fate (Daum et al. 2014; Yadav et al. 2011). In addition, WUS controls spatial distribution of *HEC1* mRNA by directly repressing its expression, which is crucial to maintain SAM integrity (Schuster et al. 2014). In turn, HEC factors form protein complexes with the bHLH transcription factor SPATULA and control the timing of stem cell differentiation by modulating phytohormonal balance (Gaillochet et al. 2017). Mechanistically, HEC factors promote cytokinin responses at the center of the SAM and repress the auxin feedback system at the periphery by transcriptionally regulating and physically interacting with AUXIN RESPONSE FACTOR 5 (ARF5) / MONOPTEROS (MP) (Figure S1; (Gaillochet et al. 2017)).

In addition to these functions, *HEC* genes control patterning in the developing fruit. *HEC* loss-of-function in *hec1,2,3* triple mutants leads to important defects in style, stigma, transmitting tract and septum formation, ultimately leading to plant sterility (Gremski et al. 2007). In this context, HEC factors directly promote the expression of the auxin-efflux carriers *PIN-FORMED 1* (*PIN1*) and *PIN3*, which pattern auxin transport and responses at the style to allow its correct development (Figure S1; (Schuster, Gaillochet & Lohmann 2015; Moubayidin & Østergaard 2014)). Importantly, *hec1,2,3* mutants also display hypersensitivity to cytokinin during style development, demonstrating the role of HEC function in balancing auxin and cytokinin responses at the gynoecium (Schuster, Gaillochet & Lohmann 2015). Similarly to HEC and SPT function, the STYLISH and NGATHA (NGA) transcription factors control style development (Martínez-Fernández et al. 2014; Eklund et al. 2010). Their activity is tightly intertwined with auxin signalling as they promote the expression of auxin biosynthesis genes including *YUCCA4*, leading to an accumulation of auxin at the apical part of the gynoecium (Martínez-Fernández et al. 2014; Eklund et al. 2010).

Given the exquisite versatility of HEC function throughout plant development, we wanted to investigate the molecular mechanisms orchestrating these multiple activities. To this end, we undertook a network approach combining protein-protein interaction screens with genome-wide profiling analyses and identified five regulatory modules associated with HEC1. Using these resources, we revealed a function for *HEC* genes in controlling flowering time and identified *NGA* genes as relevant HEC1 targets in maintaining SAM homeostasis.

## Results

### A network analysis defines HEC1 regulatory modules

The functional characterization of *HEC* genes demonstrated that they contribute to diverse developmental programs including shoot meristem activity, gynoecium patterning and photomorphogenesis (Zhu et al. 2016; Schuster et al. 2014; Schuster, Gaillochet & Lohmann 2015; Gaillochet et al. 2017). Although some regulatory interactions mediating this amazing functional diversity have been characterized, a unifying framework of the underlying mechanisms was outstanding. To fill this gap, we aimed at quantitatively identifying the core functional modules that define the diverse HEC1 activities. To this end, we first carried out two independent Yeast-Two-Hybrid screens to identify proteins with the ability to physically interact with HEC1. The first screen was done using an unbiased cDNA library derived from micro-dissected inflorescence meristems, while the other screen employed pairwise combinations against the REGIA transcription factor library (Supplementary file 1; (Castrillo et al. 2011)). Interrogating more than 1.1 10^7^ colonies in the floral library and more than 1100 individual transcription factors in the REGIA library allowed us to reliably identify 31 proteins physically interacting with HEC1 in yeast (Figure S2a-c; Supplementary file 1)(Castrillo et al. 2011). Out of these 31 hits, 12 were bHLH transcription factors, in line with the ability of basic/helix-loop-helix (bHLH) domains to mediate homo- or hetero-dimerization (Toledo-Ortiz et al. 2003). Interestingly, these factors belonged to a large spectrum of bHLH subfamilies, suggesting that HEC1 does have little or no preference to interact with specific classes of bHLH domains (Supplementary file 1). We used publicly available datasets to complement our set of HEC1 cofactors and were able to extend the list by six additional candidates. Importantly, many of the 37 factors identified were co-expressed with *HEC1*, suggesting that they could indeed functionally interact under specific developmental contexts *in vivo* (Supplementary file 1; (Klepikova et al. 2016)).

In line with previously described HEC functions, the most enriched Gene Ontology (GO) categories among HEC1 cofactors included regulation of gene expression (FDR = 7.2e-27); developmental process (FDR= 4.1e-08), response to red and far red light (FDR= 4.4e-06), shoot system development (FDR= 6.5e-04), reproductive structure development (FDR= 1.7e-02) and hormone-mediated signalling pathway (FDR= 3.41e-02) (Figure 1a)(Schuster et al. 2014; Zhu et al. 2016; Gremski et al. 2007).

**Figure 1:**
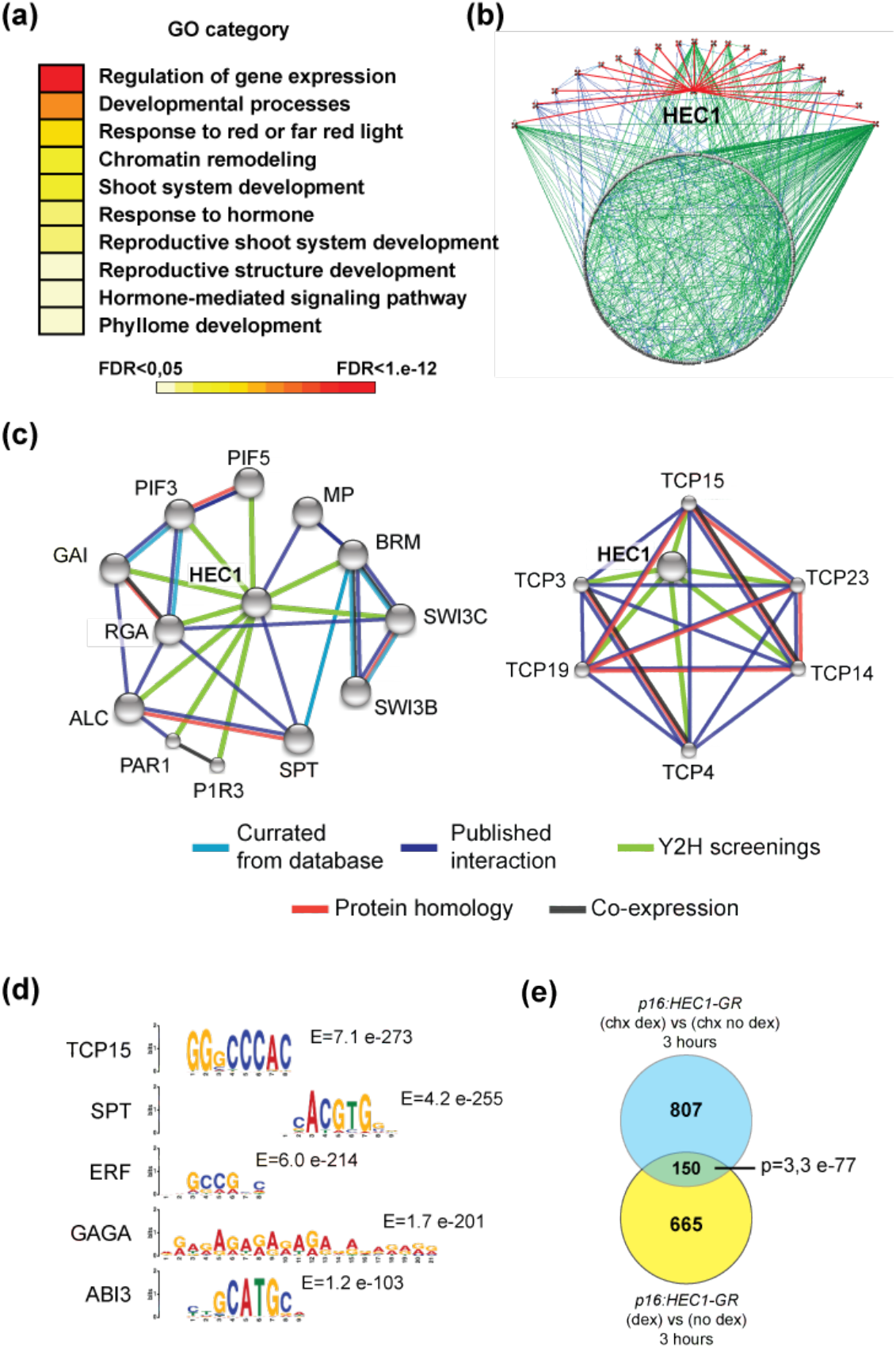
A systems analysis identifies HEC1 regulatory modules. **(a)** Over-represented gene ontology (GO) categories among identified HEC1 interactors (FDR < 0,05). **(b)** HEC1 protein-protein interaction network (interaction level 2). 20 out of 37 HEC1 cofactors identified by Yeast-Two-Hybrid assembled into a single network (Arabidopsis Interactome Mapping Consortium 2011). **(c)** HEC1 protein-protein interaction network reconstruction (interaction level 1: STRING). **(d)** Strongly enriched DNA-binding motifs representative of transcription factor families associated with HEC1-chromatin-binding regions and corresponding E-value **(e)** High confidence HEC1-early response genes as shown by overlap between *p16:HEC1-linker-GR* RNA-seq experiments 3 hours after dexamethasone (dex) treatment in the presence or absence of the protein synthesis inhibitor cycloheximide (chx). Statistical test: Benjamini-Hochberg procedure (b), hypergeometric test (e).

To elucidate the functional relationship between the 37 HEC1 interactors, we reconstructed a protein-protein interaction network, where individual nodes represent HEC1 interactors and their direct cofactors (Interaction level 2: (Arabidopsis Interactome Mapping Consortium 2011)). Interestingly, we found that 20 out of the 37 HEC1 cofactors were organized in a highly connective interaction matrix, by sharing common cofactors (Figure 1b; Supplementary file 1). This finding suggested that HEC1 may carry out its diverse functions as part of a larger regulatory unit, rather than by engaging with fully distinct complexes for each individual activity. To further zoom into this matrix and to identify interactions that could be responsible for specific HEC1 outputs, we reconstructed protein association networks only for HEC1 direct interactors using the STRING tool ((Szklarczyk et al. 2015); Supplementary file 1). The resulting interaction network again displayed high connectivity and could further be subdivided into two main sub-networks (Figure 1c; Figure S2d). The main cluster included proteins with known functions in five developmental programs: I. Light signalling as represented by PHYTOCHROME INTERACTING FACTOR 3 (PIF3), PIF5, GIBBERELLIC ACID INSENSITIVE (GAI), REPRESSOR OF GA (RGA) and PHY RAPIDLY REGULATED 1 (PAR1) (Pfeiffer et al. 2014; de Lucas et al. 2008; Feng et al. 2008; Zhou et al. 2014). II. Factors involved in the regulation of flowering time, specifically RGA, GAI and BRM (Galvao et al. 2012; Farrona et al. 2011). III. Key regulators of gynoecium development, namely SPT, ALCATRAZ (ALC), GAI and RGA (Arnaud et al. 2010; Heisler et al. 2001; Fuentes et al. 2012; Rajani & Sundaresan 2001). IV. the chromatin remodeling factors BRM, SWITCH SUBUNIT 3 (SWI3B), SWI3C (Vercruyssen et al. 2014). V. components of hormone signaling represented by MP, RGA and GAI (Hardtke & Berleth 1998; Arnaud et al. 2010). In contrast, the second cluster was characterized by weaker interactors (Figure S2a) and included exclusively TEOSINTE BRANCHED, CYCLOIDEA AND PCF (TCP) transcription factors: TCP3, TCP4, TCP14, TCP15, TCP19 and TCP23, which were shown to play divergent roles during development, including flowering transition and hormonal responses (Kubota et al. 2017; Lucero et al. 2015; Davière et al. 2014)(Table1).

The functional diversity of cofactors and the high connectivity of the reconstructed clusters suggested that HEC1 could be part of a dynamic regulatory complex, which is able to mediate distinct functions during plant development. Mechanistically, this also suggested that the physical interaction with distinct transcription factors could in turn instruct HEC1 DNA-association patterns, and thus could specify the spectrum of HEC transcriptional target genes. To further investigate this idea, we analyzed the DNA sequences of HEC1 chromatin binding regions we had previously recorded by ChIP-seq (Gaillochet et al. 2017). Indeed, HEC1 binding regions contained 227 overrepresented DNA sequences, including known target motifs of several families of transcription factors (Figure 1d; Figure S3a; (Bailey et al. 2009)). In line with our protein-protein interaction data, two of the most highly enriched sequences were the TCP-binding motif (E-value < 7.3 e-160) and the G-box (E-value < 1.4e-141) typically bound by TCP and bHLH transcription factors, respectively (Figure 1d, Supplementary file 2, (Lau et al. 2014; Pfeiffer et al. 2014)). This result not only independently supported the identification of TCP proteins as HEC1 binding partners, but also suggested that they might represent relevant output modifiers of HEC1. Furthermore, we detected an enrichment for the Auxin Response Element (ARE) (E-value < 2.6e-036) (Figure S3a; (and previously described in Gaillochet et al. 2017)), which is bound by ARFs, further supporting the functional interaction between HEC transcription factors and the auxin signalling pathway (Gaillochet et al. 2017; Boer et al. 2014). We also found a mild enrichment for the motif bound by the flowering repressor SCHLAFMUTZE (SMZ) (E-value = 1.4e-05), an AP2 transcription factor, also identified as HEC1 cofactor, but which did not group into the network of 20 connected regulators (Figure S2a, S3a). Together, these results showed that HEC1 protein-protein interaction networks correlated well with its *in vivo* DNA binding capacity, suggesting that the association with specific transcription factors could instruct the recruitment of HEC1 complexes to distinct genomic sites and in turn mediate regulatory specificity.

To follow up on this idea, we characterized the genetic circuits acting downstream of HEC function. To this end, we re-analyzed the early genome-wide transcriptional responses to HEC1 induction we had recorded in the presence or absence of the translational inhibitor cycloheximide (cyc) (Gaillochet et al. 2017). By overlapping these two datasets, we identified 150 high confidence HEC1 early response genes and defined their functional signatures by gene ontology (GO) analysis (Figure 1e, Figure S3b, Supplemental file 3). Interestingly, only two distinct categories were significantly enriched – pattern specification process (p < 9e-05), and regulation of hormonal level (p < 5.2e-05)(Figure S3b) – which were consistent with the previously described roles of *HEC* genes. Working under the hypothesis that cofactors will dictate HEC1 function by modulating DNA binding specificity and thus target gene selection, we next merged physical and genetic interactions networks to generate a putative molecular framework for HEC function (Vercruyssen et al. 2014). The resulting five core HEC1 regulatory modules supported our hypothesis, since the biological functions associated with them largely reflected the interaction network: We identified modules with annotated functions in light signaling, regulation of flowering time, gynoecium development, auxin signaling as well as an uncharacterized TCP-target network (Table 1, Figure S4). Hence, HEC1 target genes seemed to be sufficient to explain the biological functions of interaction partners and vice-versa. Since chromatin remodelers identified as class of connected HEC1 interactors do not exert their function via regulation of a small number of specific target genes, it was not surprising that we were unable to capture a core regulatory module for this functional category. Importantly, three of the modules we identified corresponded to previously described HEC functions (Zhu et al. 2016; Schuster, Gaillochet & Lohmann 2015; Gremski et al. 2007; Gaillochet et al. 2017). Furthermore, the identification of two previously undescribed regulatory modules predicted that *HEC* genes could control flowering time and also so far unknown processes together with TCP transcription factors (Table 1).

**Table 1:**
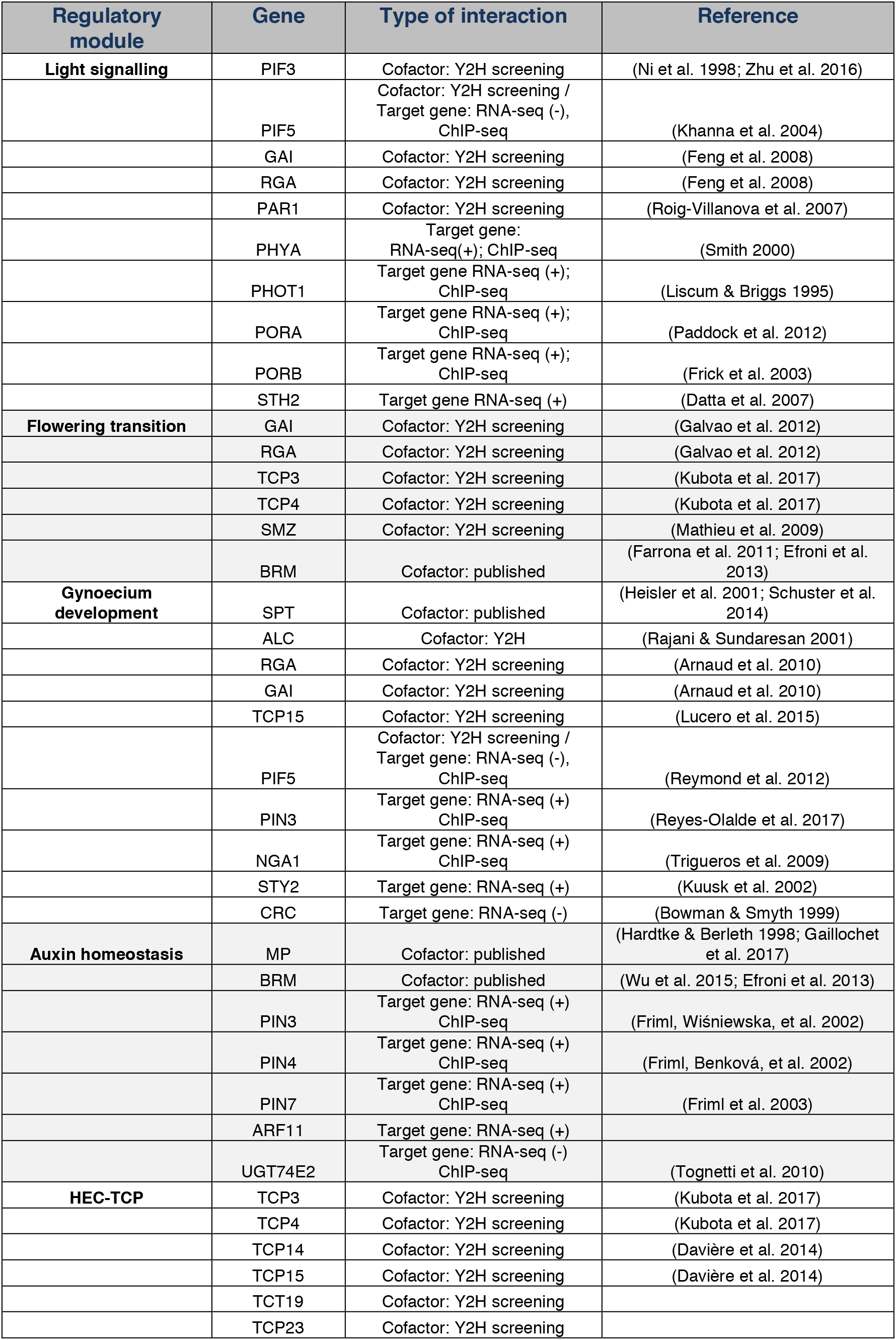
HEC1-regulatory modules. RNA-seq: elevated (+) or reduced (-) mRNA expression in response to HEC1.

### HEC activity controls flowering time

Having defined core functional modules from HEC1 regulatory networks, we next wanted to investigate their functional relevance using flowering time as a model. Together with the light signaling module, the identification of DELLA proteins, TCP3, TCP4, SMZ and BRM as HEC1 putative cofactors suggested that HEC function might also control this process (Galvao et al. 2012; Schmid et al. 2005; Kubota et al. 2017; Mathieu et al. 2009). To directly test this hypothesis, we measured flowering time of *HEC*-loss of function mutants under long-(16h light/8h darkness) and short-day (8h light/16h darkness) conditions, respectively (Figure 2a-b). Interestingly, under both light regimes, we found that among multiple *HEC* loss-of function mutants, *hec1,3* double mutants developed a reduced number of rosette leaves (Figure 2a-b, Figure S5a-d). While significant under both conditions, removing *HEC* function produced a more pronounced effect under long-days (Figure 2a-b; Figure S5c-d). Interestingly, *hec1,3* mutant flowered earlier than wild type plants under short day conditions, whereas they flowered at the same time under long day conditions, demonstrating that leaf initiation rate was modified in *hec* mutants under this light regime (Figure S5e-f).

**Figure 2:**
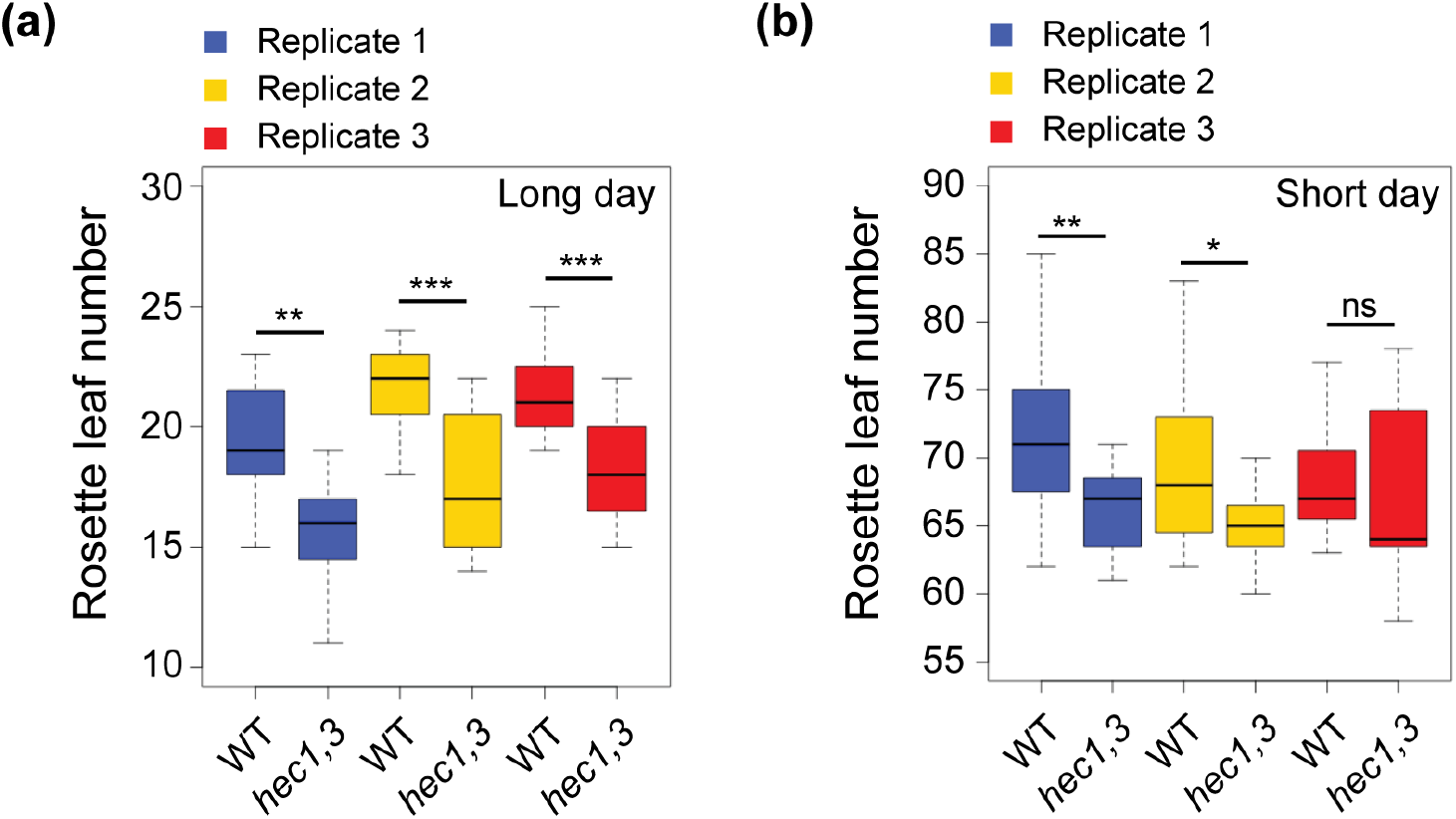
HEC function controls flowering time. **(a-b)** Flowering time assay in wild type and *hec1,3* mutants. Quantification of total rosette leaf number under long day (a) or short day conditions (b). n=15 per replicate; statistical test: Student t test: * p<0.05; ** p<0.01; *** p<0.001.

Taken together, these results revealed that *HEC* genes modulate the transition of the SAM from the vegetative to the inflorescence stage and together with the modules representing known functions, supported the biological relevance of our reconstructed network.

### Functional specificity of a HEC1 interaction module

Having uncovered a function for *HEC* genes in controlling flowering time, we next wanted to analyze the functional relationship between the individual members of a module. To this end we chose the largest regulatory module including cofactors and target genes regulating gynoecium development. In particular, we focused our analysis on SPT, ALC, RGA and GAI as putative cofactors and *NGA1* as a direct target gene (Table 1). The function of these genes during gynoecium development has thoroughly been investigated (Arnaud et al. 2010; Groszmann et al. 2011; Trigueros et al. 2009), however, their expression pattern in the shoot meristem suggested that these factors could also interact with *HEC* genes in controlling SAM development (Trigueros et al. 2009; Serrano-Mislata et al. 2017). Given the key role of HEC-SPT complex in modulating the dynamics of cell differentiation in the SAM and the high connectivity between HEC1, SPT, RGA, GAI and ALC interaction network (Figure S6a; (Schuster et al. 2014; Gaillochet et al. 2017)), we wondered whether other members of the module could have similar activities. To address this question, we first recorded ALC and RGA expression patterns at the SAM using published translational reporters (Rajani & Sundaresan 2001; Silverstone et al. 2001). We detected ALC-GUS mostly at the flower primordia and at the base of the flower petioles, whereas RGA-GFP accumulated mostly in the CZ (Figure S6b, f), suggesting that these interactors could play regionally distinct roles. To test whether these cofactors were required for HEC function in subdomains of the SAM, we analyzed the capacity of *alc* mutant or *rga gai* double mutant plants to respond to elevated HEC1 expression in the CZ or all cells of the meristem, respectively. Expression of *pCLV3:HEC1* or *p35S:HEC1* in *alc* or *rga gai* mutant backgrounds led to SAM expansion or pin-like inflorescences, respectively, which were indistinguishable from the phenotypes caused in wild-type (Figure S6c-e, S6g-i). These results demonstrated that ALC and DELLA factors were not required for HEC1 activity in the shoot meristem despite their co-expression in this tissue. Collectively, these results not only supported the idea that SPT is the predominant cofactor mediating HEC function in the SAM (Schuster et al. 2014; Gaillochet et al. 2017), but also suggested that the functional interaction between HEC1, ALC, RGA and GAI is specific to the gynoecium.

### Functional specificity of a HEC1 downstream module

Having shown that the interaction partners identified in the gynoecium module act in a tissue restricted manner, we extended our analysis to HEC1 target genes from the same module. To this end, we focused our attention on NGA transcription factors, since at the phenotypic level, *hec* and *nga* mutants exhibit related defects during gynoecium development (Schuster, Gaillochet & Lohmann 2015; Trigueros et al. 2009; Alvarez et al. 2009). From our genome-wide profiling (Gaillochet et al. 2017), *NGA1* and *NGA2* were both bound and transcriptionally activated by HEC1, suggesting that *NGA* genes could act downstream to mediate HEC function (Figure 3c-e). Importantly, HEC1 and NGA expression patterns overlapped in the gynoecium and in the SAM (Figure 3a-b; (Trigueros et al. 2009)), suggesting that their functional interaction might extend to the meristem. Consequently, to test NGA function in shoot stem cells we enhanced *NGA1* and *NGA2* expression specifically in the CZ using *pCLV3:NGA1-linker-mCherry* and *pCLV3:NGA2-linker-mCherry* (Figure 3f-l). Interestingly, about 35% of independent T1 plants displayed strong SAM fasciation, together with an over-accumulation of stem cells, just like in *pCLV3:HEC1* plants, demonstrating that *HEC* and *NGA* genes can induce similar phenotypes when their expression is elevated in the CZ (Figure 3g,j,l; (Schuster et al. 2014; Gaillochet et al. 2017)). In line with this result, the lists of genes regulated by HEC1 and NGA3 significantly overlapped, supporting a functional convergence at the transcriptional level (Figure 3m; Supplementary file 3). However, among the *pCLV3:NGA* plants, 5 to 10% displayed shoot stem cell termination and developed instead a gynoecium-like structure (Figure 3h,k). These phenotypes were never observed in HEC gain-of-function experiments (Schuster et al. 2014; Gaillochet et al. 2017), and suggested that in addition to a stem cell promoting function shared with *HEC1*, *NGA* genes also orchestrate divergent transcriptional programs.

**Figure 3:**
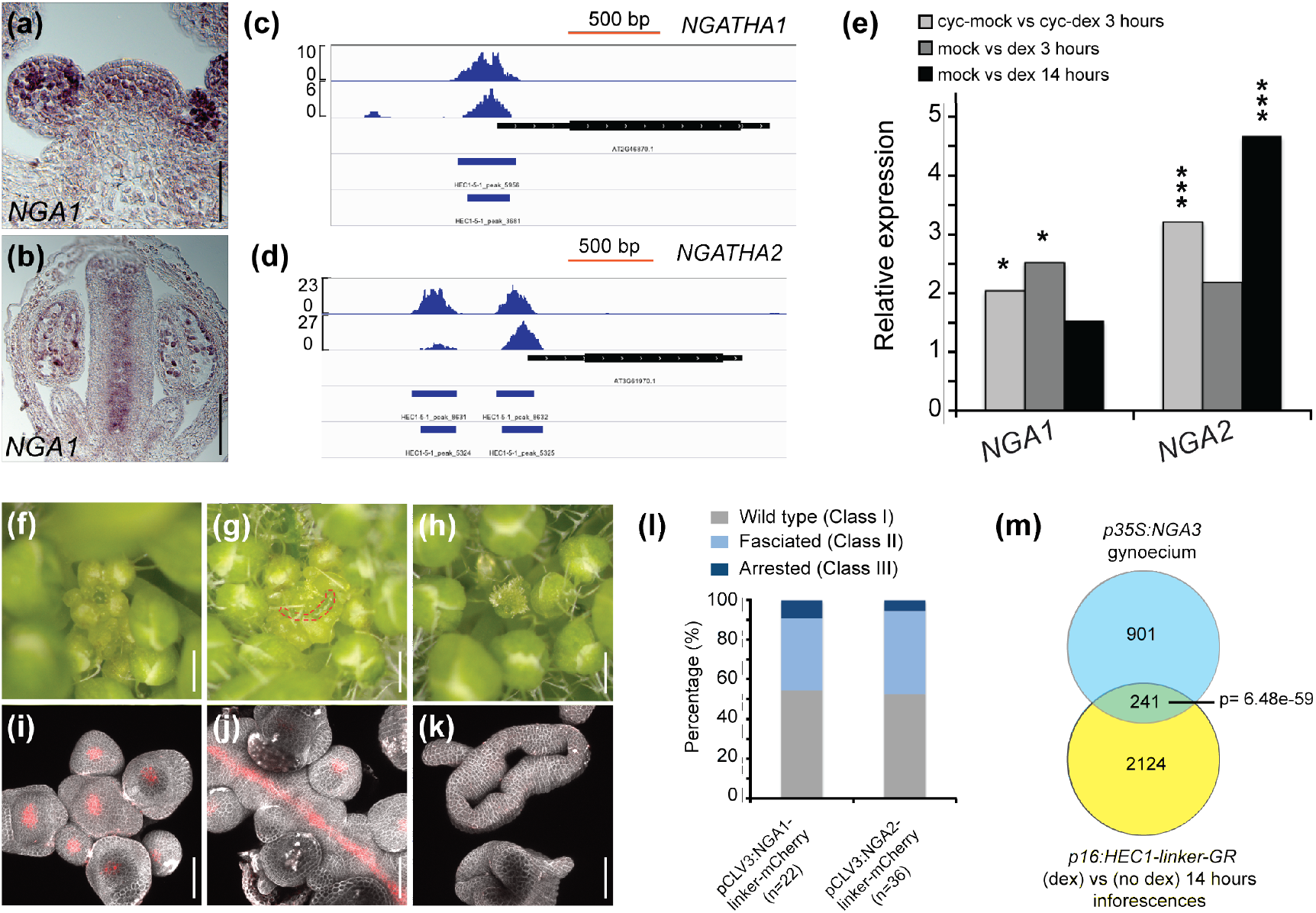
*NGA1 and NGA2* are relevant HEC1 target genes in the SAM. **(a-b)** *NGA1* expression as shown by in situ hybridization in the shoot meristem (a) or in gynoecium at stage 11-12 (b). **(c-d)** Visualization of HEC1-binding regions at *NGA1* (c) and *NGA2* (d) genomic loci. **(e)** *NGA1* and *NGA2* expression fold change 3 and 14 hours after *p16:HEC1-linker-GR* induction as measured by RNA-seq **(f-h)** Phenotypic analysis of *pCLV3:NGA1-linker-mCherry* T1 plants showing 3 classes of phenotypes: wild type-like (f), fasciated SAM (g) arrested SAM with gynoecium-like structure (h). **(i-k)** Representative view of *pCLV3:NGA2-linker-mCherry* T1 plants shoot meristems in WT-like SAM (i) fasciated SAM (j) or differentiated SAM (k). Red signal marks *NGA2-mCherry* accumulation. **(l)** Phenotypic quantification of *NGA* gain of function in the CZ. **(m)** Venn diagram showing overlap between HEC1 and NGA3-response genes. Scale bars: 50*µ*m (d-e, i-k); 500*µ*m (f-h). Statistical test: Fischeŕs exact test (EdgeR); hypergeometric test (m).

To test whether this behavior is specific to the HEC1-NGA regulatory pair, we looked for other HEC1 targets with similar function. Within the gynoecium regulatory module, we indeed found *STY2* as a high confidence positive target gene (Table 1) and *sty* mutants display gynoecium defects highly similar to *hec1,2,3,spt* (Kuusk et al. 2002). Thus, we tested whether *STY2* would also be able to control shoot stem cell activity, by expressing *STY2* specifically in stem cells using *pCLV3:STY2-linker-mCherry* and *pCLV3:mCherry-linker-STY2*. In contrast to HEC and NGA factors, activation of STY2 did not cause any SAM defects (Figure S7a), demonstrating that not all members of the HEC-gynoecium module share a hidden stem cell function.

Since *NGA* genes showed widespread expression throughout the SAM, we next wanted to test whether they could also share regulatory function with HEC at the SAM periphery. To this end, we elevated *NGA* expression specifically at the boundary region between SAM and organ primordia by a *pCUC2:NGA1-linker-mCherry* transgene. In contrast to HEC1, which blocks primordia initiation when expressed in the same setting (Gaillochet et al. 2017), NGA1 did not interfere with this process (Figure S7b-c). This suggested that *HEC* and *NGA* genes, despite their close regulatory interaction, have region specific functions: Highly convergent in the gynoecium, partially overlapping in stem cells and likely divergent at the SAM periphery. However, since so far, we had only investigated gain-of-function scenarios, we could not rule out other explanations.

Therefore, we next aimed at testing the functional dependency between HEC and NGA factors, by analyzing the capacity of HEC1 to cause phenotypes in *nga* mutants. To this end, we promoted HEC1 levels specifically in the CZ or at the boundary zone by expressing *pCLV3:HEC1-linker-GFP* and *pCUC2:HEC1-linker-GFP* in a *nga1,2* double mutant background, respectively (Figure 4). Interestingly, loss of *NGA1* and *NGA2* function did not interfere with the capacity of HEC1 to cause developmental defects at the CZ or the boundary, indicating that NGA1 and NGA2 were not required for HEC function in the SAM (Figure 4). To test whether NGA acts upstream of HEC function or that HEC and NGA act as protein complex, we conducted the reverse epistasis experiment by expressing *pCLV3:NGA1-linker-mCherry* in a *HEC* loss-of-function background (Figure S7d-e). Importantly, *HEC* genes were not required for NGA function within the stem cell region, indicating that HEC and NGA factors were able to control stem cell behavior independently from each other (Figure S7d-e). Consistently, combining HEC and NGA within the stem cells by co-expression from the *pCLV3* promoter, did not lead to synergistic effects on stem cell activity as we observed with SPT, suggesting that HEC and NGA do not have combinatorial regulatory function (Figure S5f; (Schuster et al. 2014)).

**Figure 4:**
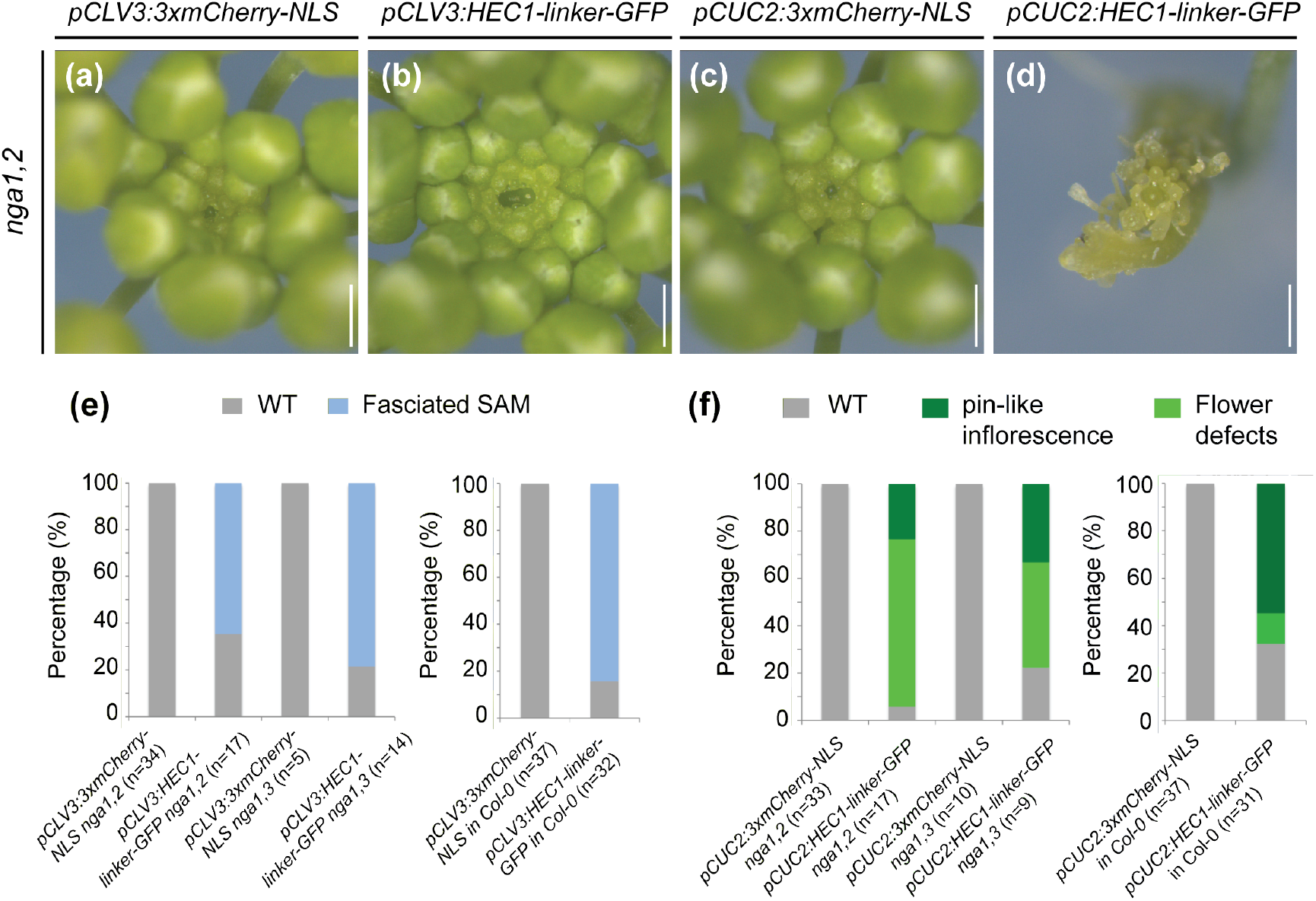
*NGA* and *HEC* genes act independently. **(a-d)** Phenotypic analysis of *nga1,2* inflorescence carrying *pCLV3:3xmCherry-NLS* (a) *pCLV3:HEC1-linker-GFP* (b) *pCUC2:3xmCherry-NLS* (c) *pCUC2:HEC1-linker-GFP* (d) in T1 plants. **(e)** Phenotypic quantification of *NGA* gain-of-function in the CZ in wild type, *nga1,2* and *nga1,3* mutant background. **(f)** Phenotypic quantification of *NGA* gain-of-function in the boundary zone (BZ) in wild type, *nga1,2* and *nga1,3* mutant background. Scale bars: 500*µ*m.

Given the strong phenotypes caused by enhancing *NGA* expression in stem cells, we finally wanted to investigate whether these factors were also required for meristem activity. Thus, we analyzed meristem size in a series of *NGA* loss-of-function mutant plants, from single to triple mutants (*nga1*; *nga1,2*; *nga1,3* and *nga1,3,4*) (Figure 5a-c; (Trigueros et al. 2009)). Surprisingly, we observed that all *nga* mutants displayed significantly larger SAMs than wild type, similar to plants with enhanced *NGA* expression (Figure 5a-c). In contrast, *hec1,2,3* had smaller shoot meristems, indicating that HEC and NGA play divergent roles in regulating SAM homeostasis despite being engaged in a close regulatory interaction (Schuster et al. 2014; Gaillochet et al. 2017). Since decreased meristem activity in *hec* triple mutants likely is caused by a reduction in cytokinin signaling (Gaillochet et al. 2017), we wanted to analyze the response to this growth-promoting hormone in *nga* mutants. To this end, we visualized cytokinin signaling output in *nga1,3* double mutant SAMs using the *pTCSn:erGFP* reporter (Zürcher et al. 2013). Consistent with their larger SAM, *nga1,3* mutants exhibited a significantly larger cytokinin signaling domain, providing a potential molecular explanation for the phenotype and demonstrating that cytokinin signalling and SAM size remained coupled in these plants (Figure 5d-f). These findings revealed that *NGA* genes were required and sufficient to modulate SAM activity and also showed that they could act independently from *HEC* factors.

**Figure 5:**
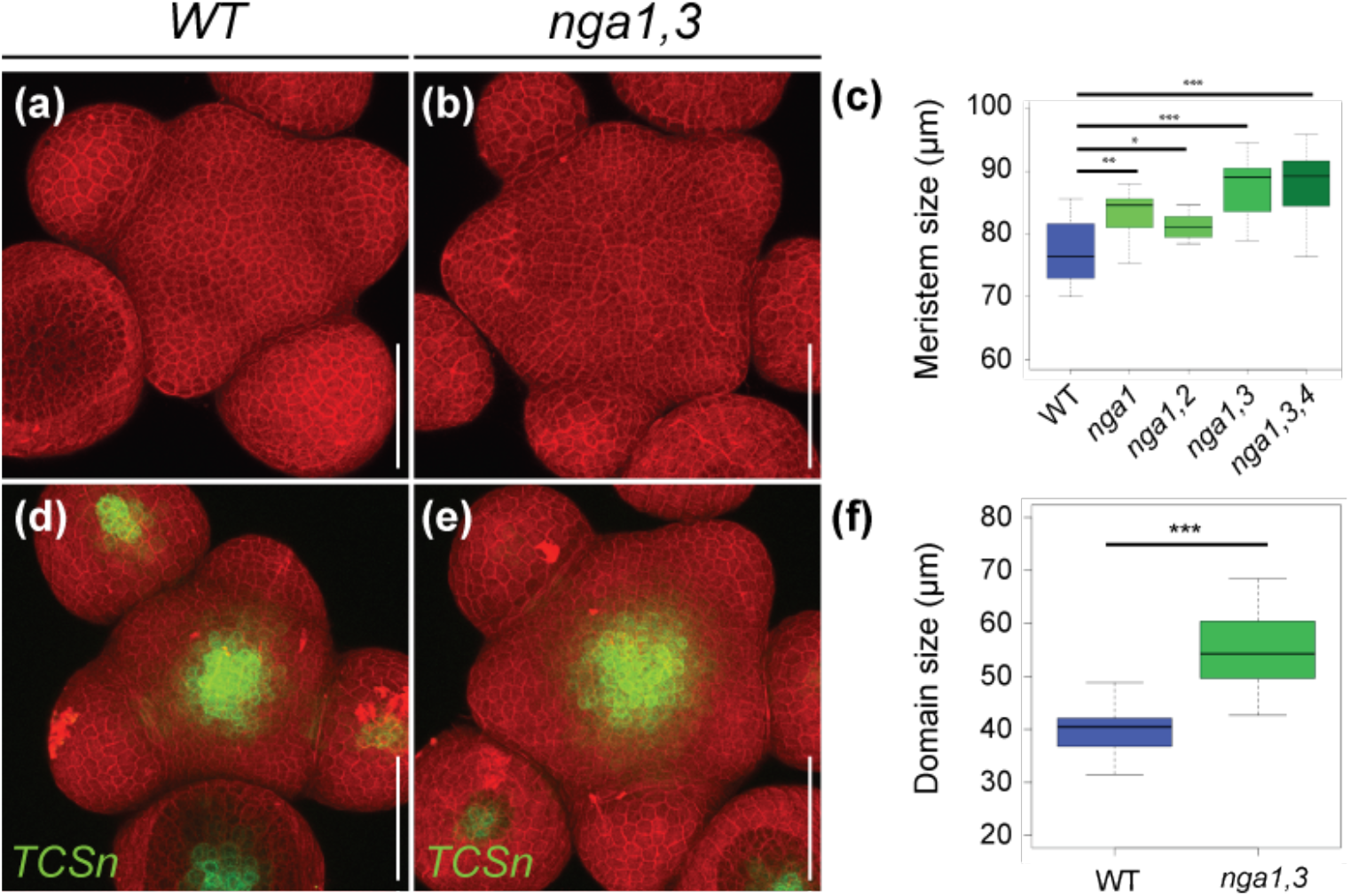
NGA function antagonizes cytokinin signalling in the SAM. **(a-b)** Representative view of wild type (a) and *nga1,3* (b) SAM. **(c)** Quantification of the shoot meristem size in wild type and *NGA* loss-of-function plants. **(d-e)** Representative view of wild type (d) and *nga1,3* (e) shoot meristems expressing the cytokinin signaling output reporter *pTCSn:erGFP*. **(f)** Quantification of cytokinin signalling domain in wild type and *nga1,3*. Scale bar: 50*µ*m.

Taken together, these results demonstrated that although *NGA* genes were directly activated by HEC1 and shared important functions in the CZ, they were not necessary for *HEC* activity in the SAM. Consistent with a role downstream of HEC1, *NGA* genes were also able to act independently of *HEC1*, if expressed appropriately. Importantly, both HEC and NGA regulators are required for proper SAM development, albeit playing opposing roles at the phenotypic and signaling level.

## Discussion

Over the course of development, plant cells respond differently to endogenous and environmental signals to adopt specific behaviors according to tissue and environmental context. However, the molecular mechanisms mediating this plasticity are currently poorly understood. Here, we used a systems-level approach combining protein-protein interaction screens and genome-wide profiling analyses to define the regulatory networks that are orchestrated by the bHLH transcription factor HEC1. From this approach, we were able to define five core regulatory modules associated with HEC function involved in the regulation of light signalling, shoot meristem activity, auxin signalling and gynoecium development, which were consistent with previous findings (Figure 6; (Zhu et al. 2016; Gremski et al. 2007; Schuster, Gaillochet & Lohmann 2015; Gaillochet et al. 2017)) and uncovered additional molecular players that could interact with *HEC* genes in these developmental contexts. It is important to note that our approach to identify interacting partners of HEC1 was limited to Yeast-Two-Hybrid screening and that additional experimental confirmation will be required to establish functionally relevant interactions *in vivo*.

**Figure 6:**
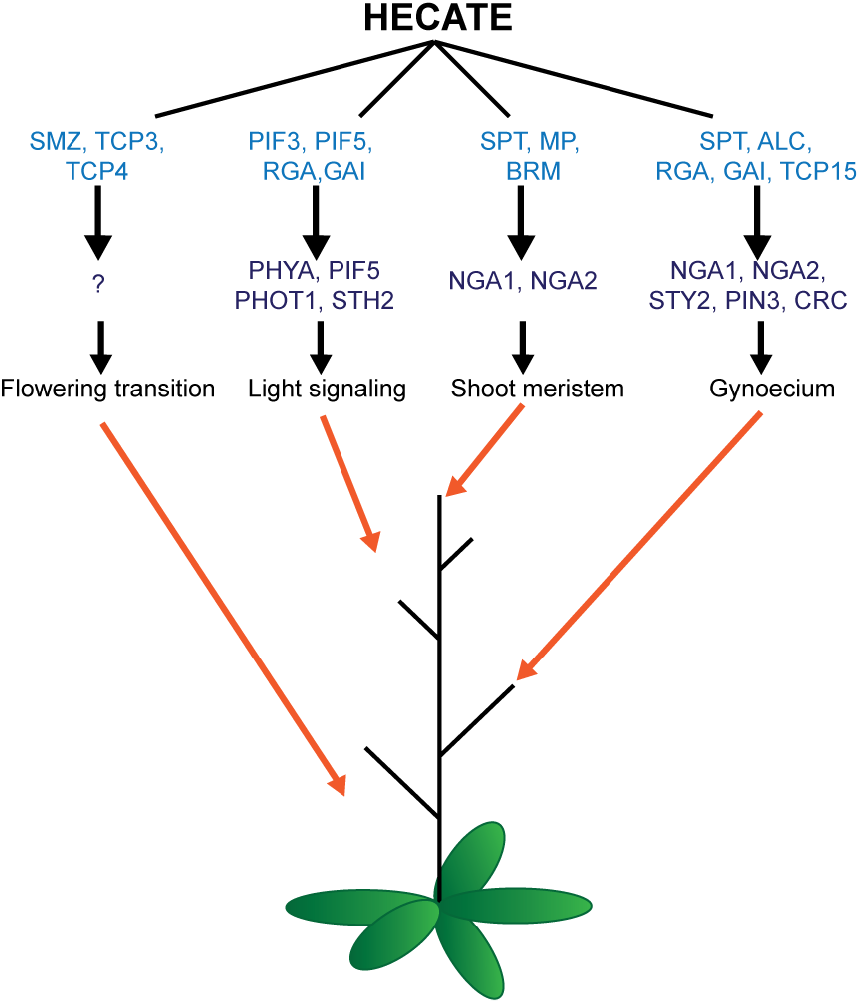
Proposed regulatory modules mediating HEC functional versatility. Genes depicted in light blue code for HEC1 cofactors whereas those in dark blue encode HEC1 direct target genes.

In addition to previously known regulatory functions, our network approach revealed that *HEC* genes modulate flowering time. Although our understanding of the molecular mechanisms underlying this specific function of HEC1 remains incomplete, its physical interaction with SMZ together with their capacity to bind similar cis-regulatory regions, including *SMZ*, *TARGET OF EAT3* (*TOE3*), *APETALA1* (*AP1*), *AP2*, *TEMPRANILLO* (*TEM1*), *FT* and *SUPRESSOR OF OVEREXPRESSION OF CONSTANS 1* (*SOC1*) (Figure S8; (Mathieu et al. 2009)), suggested that HEC1 and SMZ functionally converge to delay the transition to flowering. Furthermore, the physical interaction of HEC1 with TCP3, TCP4, DELLAs or BRM may also play a role during this process. Thus, it will be crucial in future studies to further characterize HEC1 transcriptional activity during flowering transition in order to assess the functional convergence with these putative cofactors. An alternative explanation for HEC1’s role in flowering time could be the modulation of light signalling (Zhu et al. 2016) and it will therefore be important to devise strategies to experimentally determine the relative contribution of these pathways.

Intriguingly, we identified a regulatory module composed of multiple TCP transcription factors, which are frequently found in interaction screens, and thus may represent false positives. However, recent large-scale studies also suggest that particular TCPs can constitute hubs in protein-interaction networks and hence, a large number of diverse interactions are also expected for these proteins (reviewed in (Bemer et al. 2016)). Our finding that DNA binding motifs for TCP factors are highly enriched in HEC1 chromatin interaction domains, lends support to the idea that TCPs indeed are relevant to tune HEC activity. To clarify this point, it will be important to further investigate to what extent their physical interaction with HEC1 is functionally relevant under multiple developmental contexts.

We found that many putative cofactors were shared between functional modules, suggesting that multiple developmental processes might converge at these regulatory nodes. This finding could suggest that HEC1 acts as part of a dynamic signaling complex, rather than through interaction with individual binding partners. Notably, DELLA proteins are key repressors of gibberellic acid signalling and were additionally shown to control light signalling, flowering transition, gynoecium development and inflorescence meristem size (Feng et al. 2008; Arnaud et al. 2010; Galvao et al. 2012; Serrano-Mislata et al. 2017). The role of DELLA proteins as a regulatory hub and their physical interaction with HEC1 suggests that HEC1 could function at this interface by directly modulating their activity. It will thus be exciting to further dissect these interactions under multiple developmental contexts.

In addition, using the HEC1 regulatory network as a springboard, we investigated so far undescribed roles of putative HEC1 cofactors or direct target genes in the regulation of the shoot apical meristem. We found that the physical interaction between HEC1 and ALC or DELLAs is not required for HEC function in the SAM. Rather, the previously described role of ALC and DELLAs during gynoecium development suggests that the protein complex may play a role specifically in this context (Groszmann et al. 2008; Arnaud et al. 2010). Furthermore, the high connectivity of the protein-protein interaction network between HEC1, SPT, ALC and DELLAs supported the idea that these factors may functionally interact in a larger protein complex.

We also found that *NGA1* and *NGA2* were directly activated by HEC1 at the transcriptional level. Despite the fact that NGA and HEC factors display common transcriptional signatures and that they can both promote stem cell domain and SAM expansion when expressed from the *CLV3* promoter, epistasis analysis revealed that these factors likely control SAM activity independently. Mechanistically, these results suggest that their regulatory activity converges at common nodes to control the expression of key target genes. However, in contrast to *hec1,2,3*, all *nga* mutant analyzed displayed larger SAMs, suggesting that *NGA* genes have divergent functions in controlling meristem and stem cell activity. This discrepancy between gain and loss-of-function phenotypes could result from different roles played by NGA factors in space and time within the shoot meristem as in the case of HEC1 (Schuster et al. 2014; Gaillochet et al. 2017). Alternatively, the elevation of *NGA* expression in stem cells may trigger so far unknown feedback loops that overcompensate for enhanced NGA mRNA levels. Thus, to further clarify NGA function and the regulatory circuit underpinning their activity in shoot stem cells, it will be important to precisely resolve their spatio-temporal activity within the SAM and to identify some of the target genes mediating their function.

The functional and molecular convergence of HEC and NGA pathways in the gynoecium, suggest that their regulatory interaction in this tissue may vary from the SAM (Trigueros et al. 2009; Schuster, Gaillochet & Lohmann 2015; Martínez-Fernández et al. 2014). This is in line with our previous findings where we proposed that HEC regulatory function in the SAM and in the gynoecium might diverge (Schuster, Gaillochet & Lohmann 2015). However, our understanding of the mechanisms mediating HEC function in these contexts still remain incomplete and thus, it will also be important to comparatively investigate HEC regulatory interactions in the SAM and in the gynoecium, for example by profiling HEC1 DNA-binding pattern and by recording transcriptional responses after HEC1 induction in individual tissues or developmental stages.

Taken together, we identified a set of regulatory modules that could encode for HEC functional versatility observed across plant development (Figure 6). In this model, we propose that HEC factors can form multiple protein complexes that in turn could guide their association to distinct DNA regions in order to differentially regulate gene expression and trigger specific developmental programs. Along the same idea, the animal transcription factor SEX DETERMINING REGION Y-Box 2 (SOX2) physically associates with multiple protein partners, which in turn defines the DNA-binding affinities of the complexes and the range of genes transcriptionally regulated, ultimately encoding multiple regulatory functions from maintenance of embryonic stem cell pluripotency to neural stem cells differentiation ((Adachi et al. 2013); reviewed in (Kondoh & Kamachi 2010)). In parallel to this scenario, multiple processes such as protein dosage, differential chromatin accessibility leading to differential cellular responsiveness or domain-specific post-translational modification could also participate in instructing HEC functional specificity. Thus, it will be important to assess the contribution of these individual regulatory processes and to further characterize how they may influence HEC molecular functions.

## Material and methods

### Cloning

*pCLV3:HEC1-linker-GFP, pCUC2:HEC1-linker-GFP, pCLV3:mCherry-linker-NGA1, pCLV3:mCherry-linker-NGA2, pCLV3:NGA1-linker-mCherry, pCLV3:NGA2-linker-mCherry, pCUC2:NGA1-linker-mCherry, pCLV3:3xmCherry-NLS, pCUC2:3xmCherry-NLS* were generated using Green Gate cloning system (Lampropoulos et al. 2013). *pCLV3:HEC1* and *p35S:HEC1* were previously published (Schuster et al. 2014).

*pGBKT7-HEC1* was generated by ligating *HEC1* CDS with EcoRI and PstI overhangs in pGBKT7 vector. *pGADT7-ALC* were generated by ligating *SPT* and *ALC* CDS with EcoRI and BamHI overhangs in pGADT7 vector. *pDEST22-HEC1* was generated by recombination into pDONOR221 and pGreenIIS destination vectors using the Gateway cloning system (Thermo Fisher Scientific; Waltham, USA).

For pGGC-NGA1 and pGGC-NGA2, NGA1 and NGA2 cds were amplified with primers carrying T7 or SP6 sites and further ligated in pGGC plasmids using Eco31I overhangs (Lampropoulos et al. 2013).

### Transgenic lines

Transgenic plant lines were generated using standard floral dipping protocols (Clough & Bent 1998). *Agrobacterium tumefasciens* strain ASE was used.

### Plant material

*hec1,3* (Gremski et al. 2007)*; hec1,2,3* (Schuster et al. 2014)*; nga1; nga1,2; nga1,3; nga1,3,4* (Trigueros et al. 2009); *gai-t6 rga-24* (Silverstone et al. 1998; Peng et al. 1997); *pTCSn:erGFP* (Zürcher et al. 2013), *pCLV3:HEC1-linker-GR*, *p16:HEC1-linker-GR* (Gaillochet et al. 2017), *pALC:ALC-GUS* (Rajani & Sundaresan 2001), *pRGA:GFP-RGA* (Silverstone et al. 2001) were previously described. *alcatraz* mutant line corresponds to the insertion line SALK_103763 (Alonso et al. 2003)

### Plant growth, treatments and phenotype analysis

Plants were grown at 23°C, 65% humidity under long day conditions (16h light/8h dark) under LED lights (Philips, Amsterdam, Netherlands). Plant phenotypes were assessed 30 days after germination.

For flowering time assays, plants were grown at 22°C, 70% humidity on Rockwool blocks and watered with 1mg/l HYPONeX nutrient solution (ScottsMiracle-Gro, Mraysville, US). Plants were exposed to 100 *µ*mol m-2s-1 light (LED lights) under long day (16h light / 8h dark) or in short day (8h light / 16h dark) cycles. Plant positions within trays were randomized to reduce biases from environmental cues on flowering time. Total rosette leaf number was quantified in biological triplicates by measuring plants that were sown out at the same time point and grown at different locations of the growth room.

### Histology

For GUS staining, plant tissues were harvested in 90% aceton, prefixed for 20 minutes and washed with cold staining buffer without X-Gluc. Tissues were next transferred to staining buffer and incubated at 37°C in the dark. GUS signal was regularly checked and stopped before over-staining by transferring tissues in 70% EtOH. Embedding and sectioning was performed as described in (Medzihradszky et al. 2014).

In situ hybridization experiments were performed as previously described in (Medzihradszky et al. 2014).

### Yeast two hybrid assays and screenings

For small-scale yeast-two-hybrid assay, bait construct pGBKT7-HEC1 was transformed in Ah109 yeast strain as described in (Gietz & Schiestl 2007) and tested for auto-activation by mating with Y187 strain carrying empty pGADT7 vector. Diploid colonies were re-suspended in water and equal volumes were dropped on -L,W selective minimal medium to test for colonies growth and -H,L,W with an increasing concentration of 3-Amino-1,2,4-triazole (3-AT) to test for the degree of auto-activation. To test HEC1-ALC interaction, HEC1 bait strain was mated with ALC prey strain (pGADT7-SPT in Y187) and selected on –H,L,W supplemented with 20 mM 3AT.

For the unbiased yeast-two-hybrid cDNA library screening, the HEC1 bait strain was mated with a prey strain carrying a floral cDNA library, which was prepared as described in Matchmaker^TM^ gold yeast two-hybrid system user manual (Clontech; Mountain View; USA). Diploid colonies containing a potential HEC1 interactor were selected on –H,L,W medium supplemented with 15mM 3AT, isolated, resuspended in water and dropped on –H,L,W + 15mM 3AT medium for confirmation. DNA was extracted according to a modified version of the Smash and Grab protocol (Hoffman & Winston 1987). Cell pellets were resuspended in breaking buffer, phenol/chloroform/isoamyl solution with glass beads and vigorously vortexed for 1 to 2 minutes. Aqueous phase was next transferred to a new tube and DNA was precipitated with 100% ethanol. After centrifugation DNA pellet was washed with 70% ethanol, air-dried and re-suspended in TE buffer. Isolated plasmids were used as template for PCR amplification and further identification of the candidate cofactor by DNA sequencing (Eurofins genomics, Les Ulis, France).

For REGIA library screening, HEC1 bait strain was generated by transforming pDEST32-HEC1 in pJ69-4α yeast strain. Bait strain was transformed with pDEST22 vector and selected on –H,L,W media supplemented with 3-AT to test for auto-activation. Next, pJ69-4α strain was mated with pJ69-4A prey stain containing the REGIA library (Castrillo et al. 2011). Diploid colonies were selected on –A,L,W medium and on –H,L,W supplemented with 1mM and 5mM 3-AT as described in (de Folter & Immink 2011).

### Image acquisition and analysis

Confocal images were acquired on Nikon A1 Confocal with a CFI Apo LWD 25x water immersion objective (Minato, Tokyo, Japan). Image processing and analysis was performed as described in (Gaillochet et al. 2017).

Binocular pictures were recorded using digital Camera AxioCam HRC (Carl Zeiss; Oberkochen, Germany).

### Bioinformatic analysis

HEC transcriptional and genome-wide DNA binding profiles were previously published, and conducted as described in (Gaillochet et al. 2017). Motif enrichment analysis was performed as described in (Gaillochet et al. 2017), using JASPAR 2016 as motif input. Only binding motifs representative for transcription factor families were displayed. A complete list of binding motifs can be found in supplementary file 2.

Gene ontology (GO) enrichment analysis among high confidence HEC1 response genes was performed with AgriGO (Du et al. 2010).

Gene regulatory modules were reconstructed by overlapping protein-protein interaction data with high confidence HEC1-response gene lists. Gene function was determined by mining published literature.

NGATHA3-response genes were obtained from (Martínez-Fernández et al. 2014).

HEC1 protein-protein interaction network with interaction level 2 was generated using VisANT on the Arabidopsis interactome online tool (Hu et al. 2004; Arabidopsis Interactome Mapping Consortium 2011).

First-level PPI clusters and GO analysis of HEC1 protein-protein interaction network were constructed using the STRING tool (Mering 2004).

### Primers

A detailed list of primers used in this study can be found in supplementary file 4

## Author contributions

CG performed all experiments at the exception of the Yeast-Two-Hybrid screening using the transcription factor library conducted by FW and RI and the flowering time assay performed by SJ, RI and GA.

CG and JL: designed experiments, analyzed data and wrote the manuscript

## Acknowledgements

We thank Cristina Ferrandiz for sharing *nga* mutant seeds, Venkatesan Sundaresan for sharing *pALC:ALC-GUS* lines, Claus Schwechheimer for *gai-t6 rga-24* line and all the members of the laboratory for critically reading the manuscript.

---

**Figure S1:**
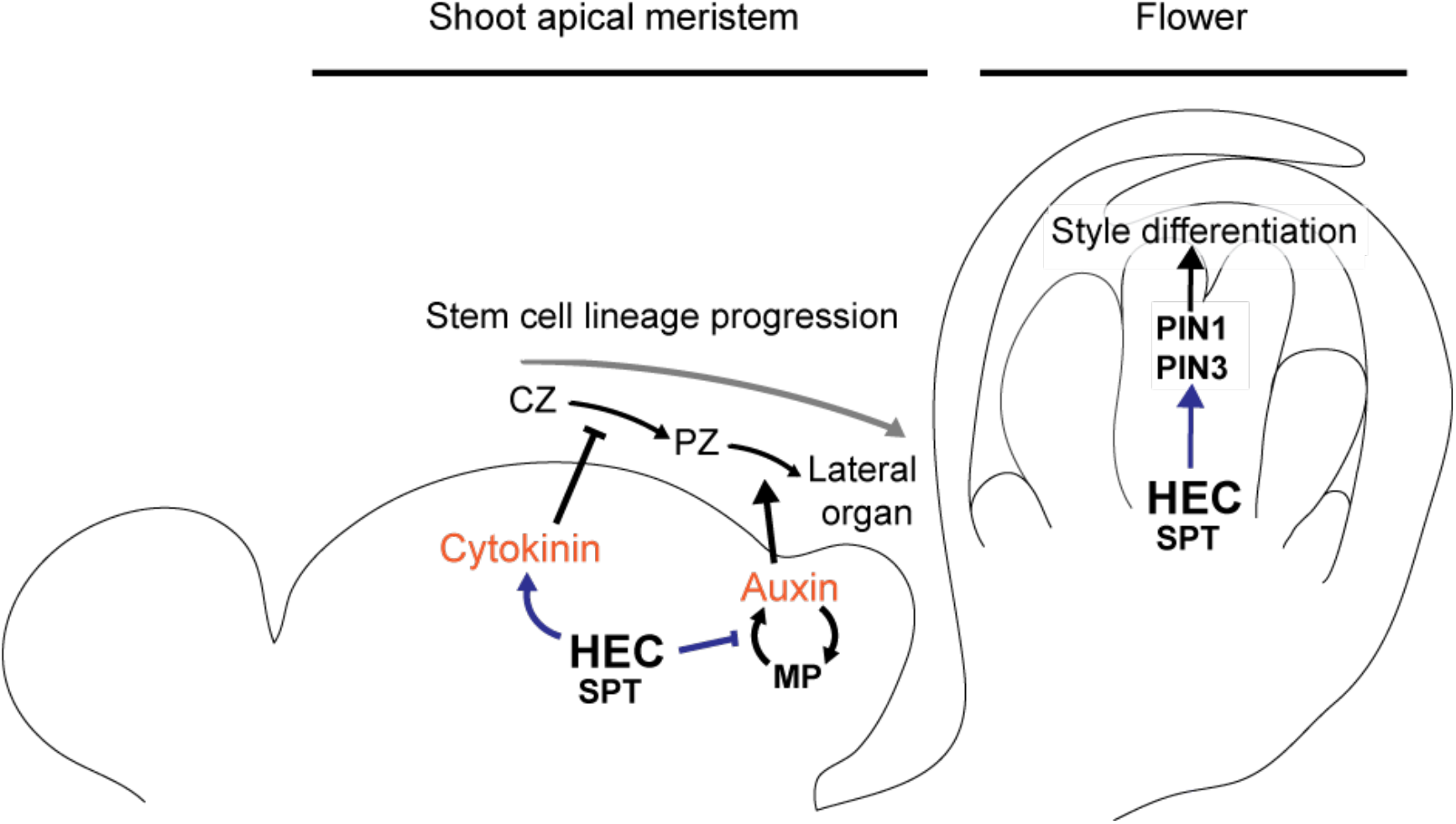
Schematic depicting HEC regulatory function in the shoot meristem and in the gynoecium.

**Figure S2:**
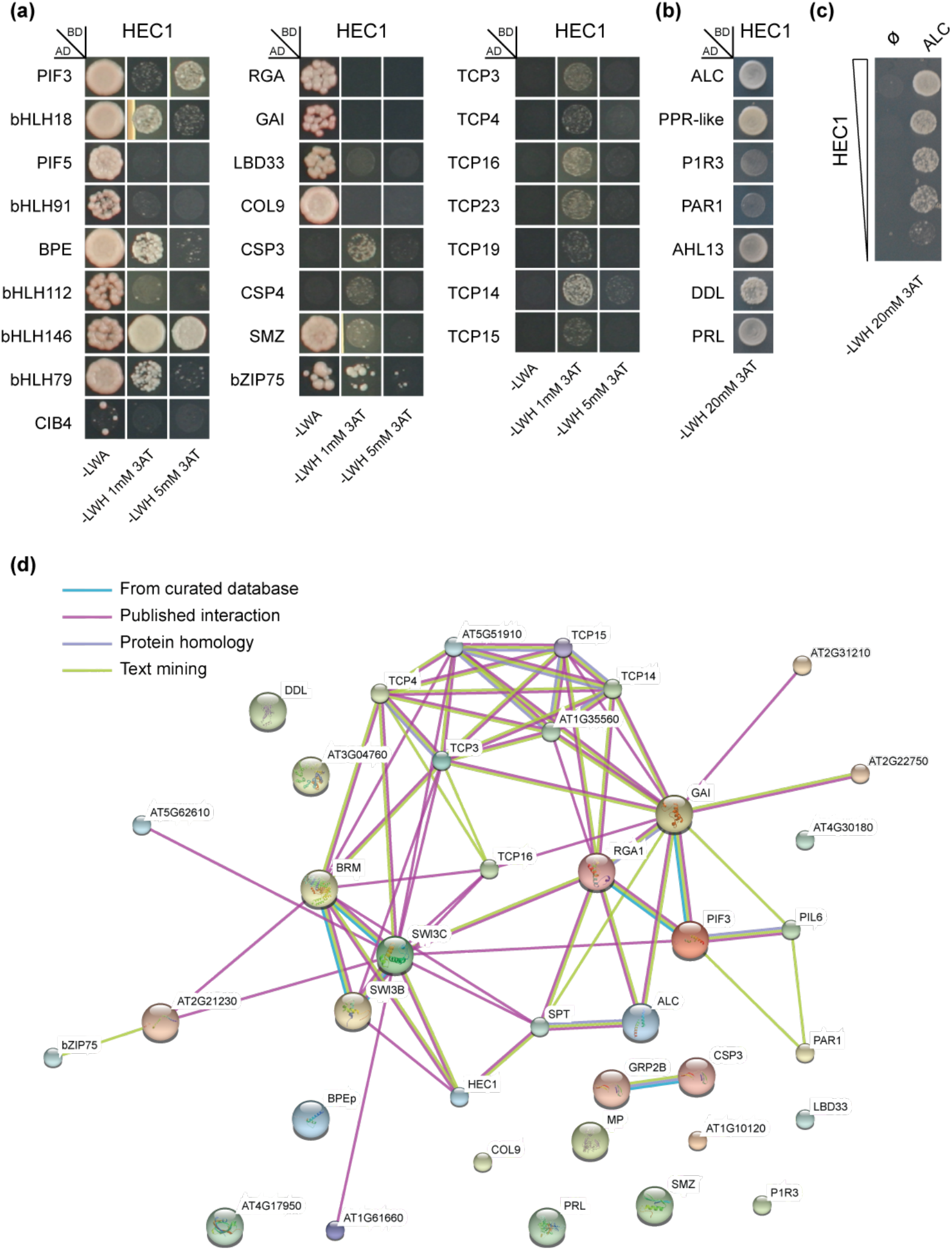
**(a)** Positive clones obtained from Yeast-Two-Hybrid screening with the REGIA transcription factor library and replicated on 3 different selective media **(b)** Positive clones obtained from Yeast-Two-Hybrid using a floral cDNA library and spotted on a selective medium. **(c)** Small scale Yeast-Two-Hybrid assay testing physical interaction between HEC1 and ALCATRAZ. Clones were plated using gradual dilution. **(d)** Interaction network output from STRING tool.

**Figure S3:**
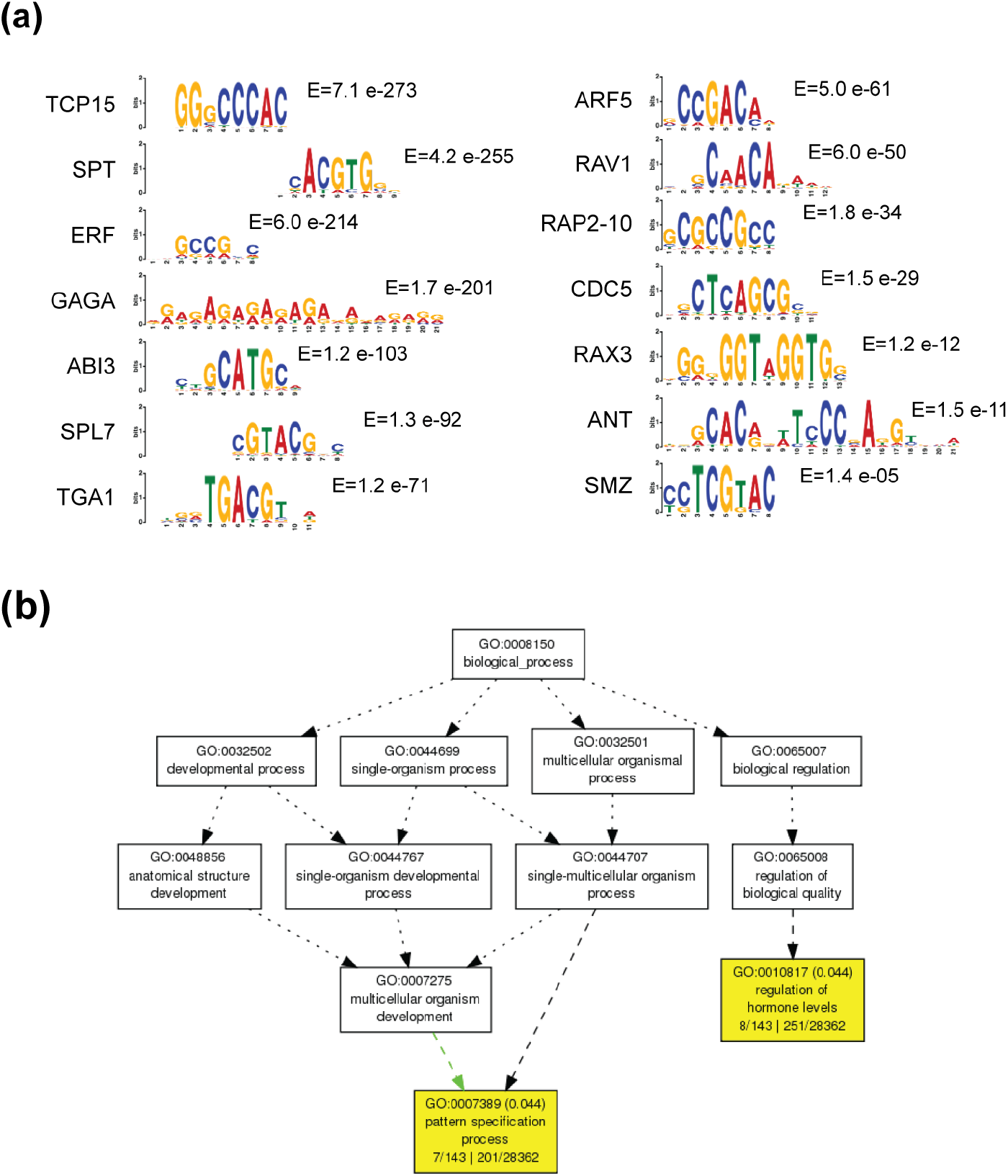
(a) DNA motifs representative of various transcription factor family enriched under HEC1-bound regions in ChIP-seq experiment, together with corresponding E-value (b) Enriched GO categories among high confidence HEC1-early response genes (from inflorescence RNA-seq; FDR < 0,05).

**Figure S4:**
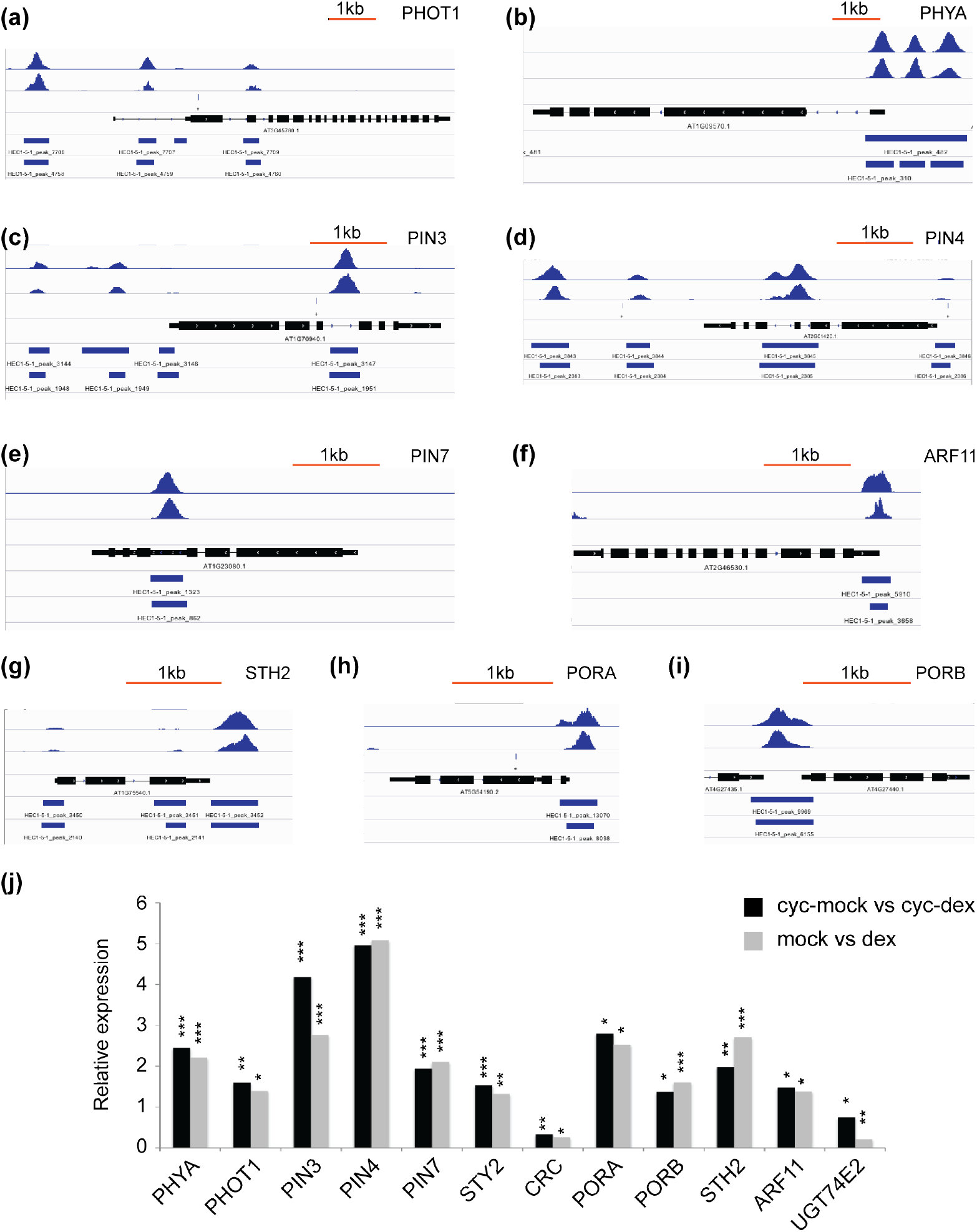
**(a-i)** Visualization of HEC1 DNA-binding pattern based on ChIP-seq experiments at the genomic loci of PHOT1 (a), PHYA (b), PIN3 (c), PIN4 (d), PIN7 (e), ARF11 (f), STH2 (g), PORA (h) and PORB (i). **(j)** Relative gene expression after *p16:HEC1-linker-GR* induction with dexamethasone, supplemented or not with cycloheximide. Statistical test: Fischeŕs exact test (EdgeR), * p<0.05, ** p<0.01, *** p<0.001 (f).

**Figure S5:**
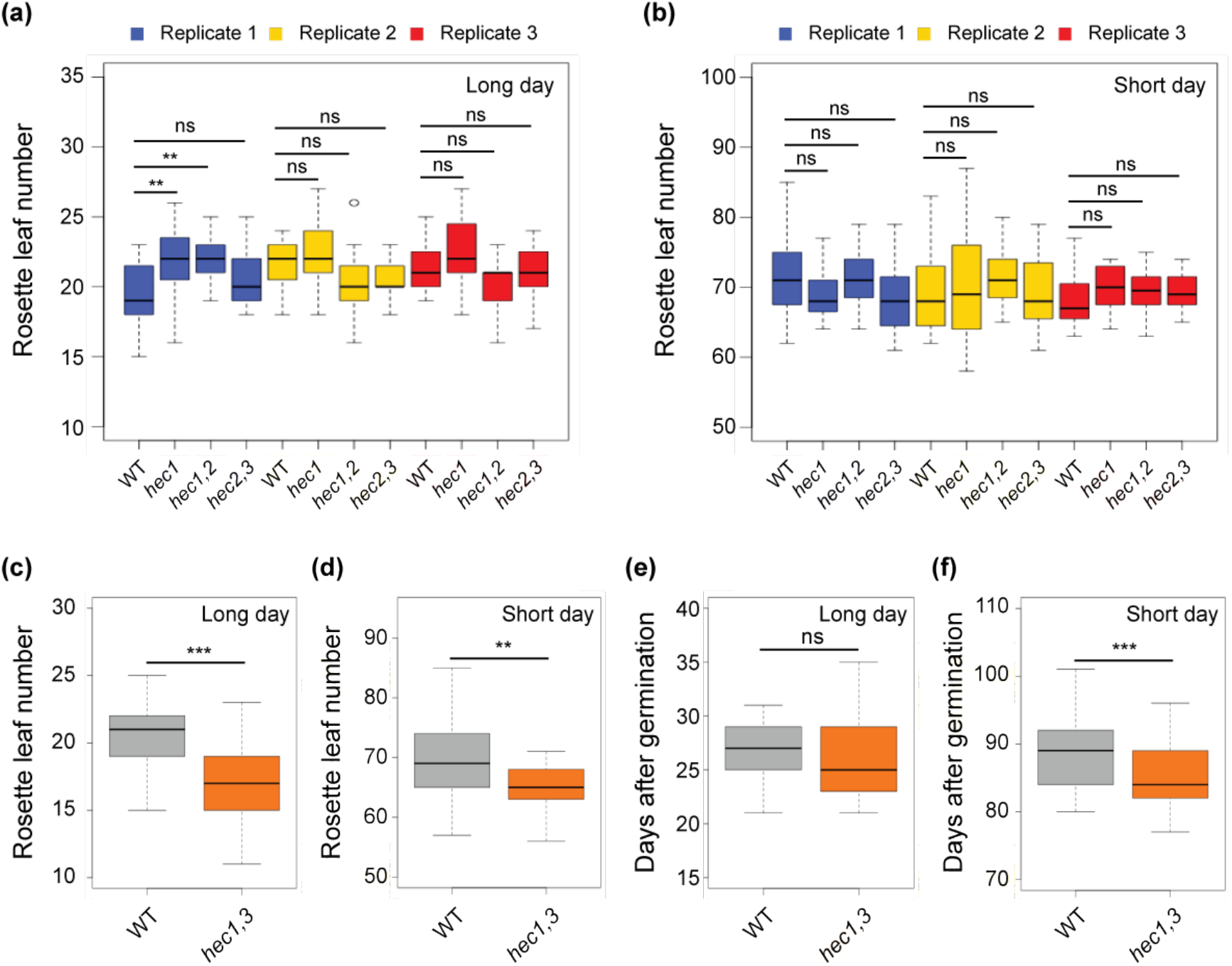
**(a-b)** Flowering time assay in wild type, *hec1*, *hec1,2* and *hec2,3* mutants (n=15 per replicate). Quantification of total rosette leaf number under long day (a) or short day conditions (b). **(c-f)** Flowering time assay in pooled wild type and *hec1,3* mutants (n=45). Quantification of total rosette leaf number (c-d) and number of days to bolting (e-f) under long day (c,e) or short day (d,f) conditions. Statistical test: Student t test: * p<0.05; ** p<0.01; *** p<0.001.

**Figure S6:**
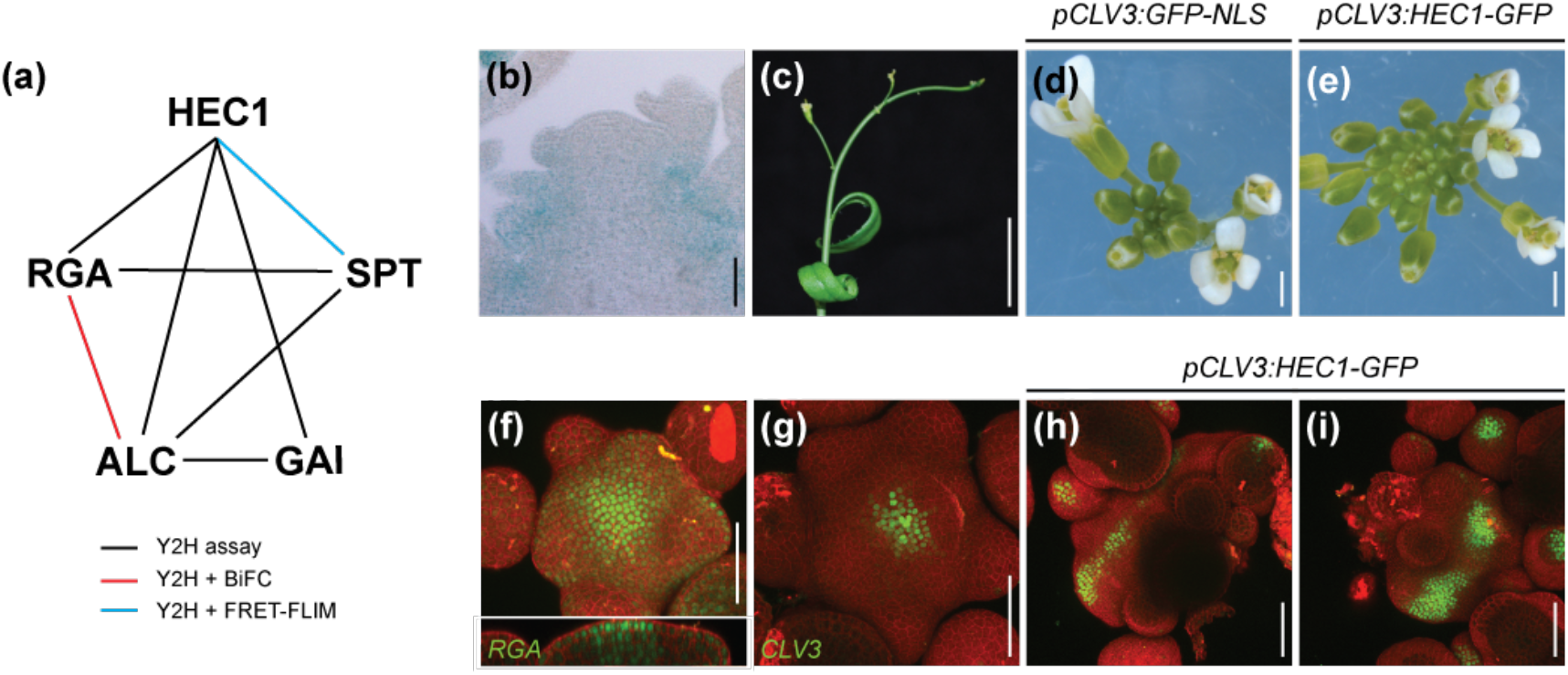
**(a)** Protein-protein interaction network between HEC1, SPT, ALC, RGA and GAI proteins. **(b)** GUS staining showing *pALC:ALC-GUS* expression in a SAM transversal cross-section. **(c-e)** *alcatraz* inforescence expressing *p35S:HEC1* (c), *pCLV3:GFP-NLS* (d), *pCLV3:HEC1-linker-GFP* (e). **(f)** Wild type meristem expressing *pRGA:GFP-RGA*. **(g-i)** Wild type (g,h) and *rga,gai* (i) meristems expressing *pCLV3:GFP-NLS* (g) and *pCLV3:HEC1-linker-GFP* (h-i). Scale bars: 50*µ*m (b,f-i); 1mm (d-e); 1cm (c).

**Figure S7:**
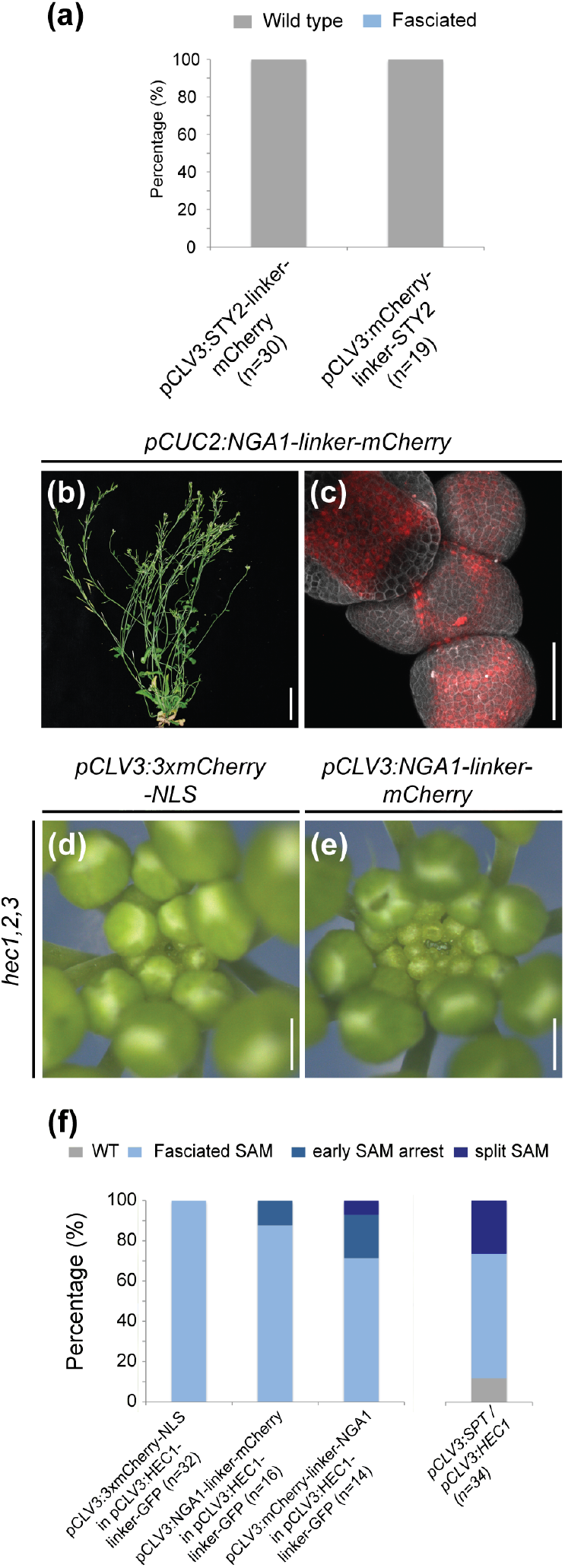
**(a)** Phenotypic quantification of inflorescence meristems expression *pCLV3:STY2-linker-mCherry* or *pCLV3:mCherry-linker-STY2*. **(b-c)** T1 plant expressing *pCUC2:NGA1-linker-mCherry.* Plant aboveground architecture (b), representative view of the SAM (c) Red signal depicts NGA1-linker-mCherry accumulation. **(d-e)** *hec1,2,3* mutant inflorescences expressing *pCLV3:mCherry-NLS* (d)(n=2) or *pCLV3:NGA-linker-mCherry* (e)(n=2). **(f)** Phenotypic quantification of T1 plants co-expressing NGA1 or SPT (Schuster et al. 2014) with HEC1 in the CZ. Scale bars: 50*µ*m (c); 500*µ*m (D-E); 5cm (b).

**Figure S8:**
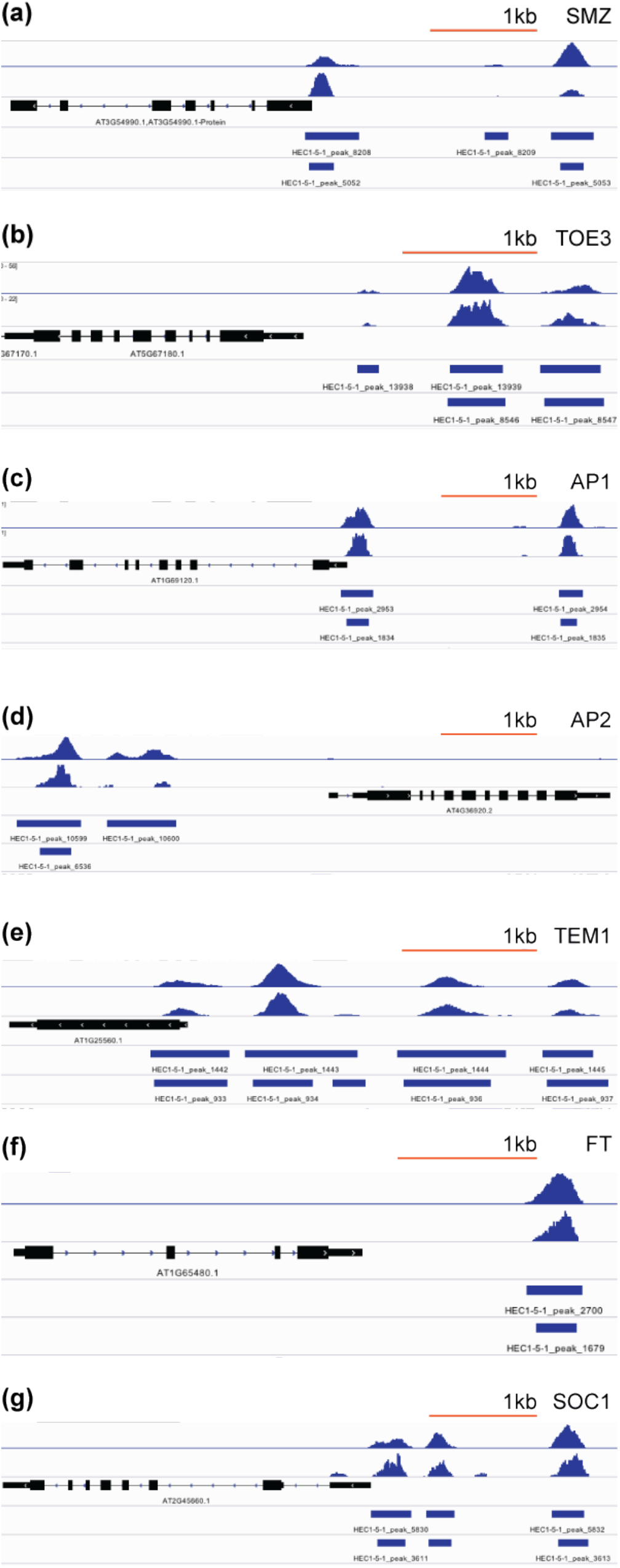
**(a-i)** Visualization of HEC1 DNA-binding pattern based on ChIP-seq experiments at the genomic loci of *SMZ* (a), *TOE3* (b), *AP1* (c), *AP2* (d), *TEM1* (e), *FT* (f) and *SOC1* (g).

## References

Adachi, K. et al., 2013. Context-Dependent Wiring of Sox2 Regulatory Networks for Self-Renewal of Embryonic and Trophoblast Stem Cells. Molecular Cell, 52(3), pp.380–392.

Alonso, J.M. et al., 2003. Genome-wide insertional mutagenesis of Arabidopsis thaliana. Science, 301(5633), pp.653–657.

Alvarez, J.P. et al., 2009. The NGATHA distal organ development genes are essential for style specification in Arabidopsis. The Plant cell, 21(5), pp.1373–1393.

Arabidopsis Interactome Mapping Consortium, 2011. Evidence for network evolution in an Arabidopsis interactome map. Science, 333(6042), pp.601–607.

Arnaud, N. et al., 2010. Gibberellins control fruit patterning in Arabidopsis thaliana. Genes & Development, 24(19), pp.2127–2132.

Bailey, T.L. et al., 2009. MEME SUITE: tools for motif discovery and searching. Nucleic Acids Research, 37, pp.W202–W208.

Bäurle, I. & Dean, C., 2006. The Timing of Developmental Transitions in Plants. Cell, 125(4), pp.655–664.

Bemer, M. et al., 2016. Cross-Family Transcription Factor Interactions: An Additional Layer of Gene Regulation. Trends in Plant Science, pp.1–15.

Boer, D.R. et al., 2014. Structural basis for DNA binding specificity by the auxin-dependent ARF transcription factors. Cell, 156(3), pp.577–589.

Bowman, J.L. & Smyth, D.R., 1999. CRABS CLAW, a gene that regulates carpel and nectary development in Arabidopsis, encodes a novel protein with zinc finger and helix-loop-helix domains. Development, 126(11), pp.2387–2396.

Brand, U. et al., 2000. Dependence of stem cell fate in Arabidopsis on a feedback loop regulated by CLV3 activity. Science, 289(5479), pp.617–619.

Castrillo, G. et al., 2011. Speeding Cis-Trans Regulation Discovery by Phylogenomic Analyses Coupled with Screenings of an Arrayed Library of Arabidopsis Transcription Factors M. A. Blázquez, ed. PLoS ONE, 6(6), p.e21524.

Clough, S.J. & Bent, A.F., 1998. Floral dip: a simplified method for Agrobacterium-mediated transformation of Arabidopsis thaliana. The Plant journal, 16(6), pp.735–743.

Datta, S. et al., 2007. SALT TOLERANCE HOMOLOG2, a B-box protein in Arabidopsis that activates transcription and positively regulates light-mediated development. The Plant cell, 19(10), pp.3242–3255.

Daum, G. et al., 2014. A mechanistic framework for noncell autonomous stem cell induction in Arabidopsis. Proceedings of the National Academy of Sciences, 111(40), pp.14619–14624.

Davière, J.-M. et al., 2014. Class I TCP-DELLA interactions in inflorescence shoot apex determine plant height. Current biology : CB, 24(16), pp.1923–1928.

de Folter, S. & Immink, R.G.H., 2011. Yeast protein-protein interaction assays and screens. Methods in molecular biology (Clifton, N.J.), 754, pp.145–165.

de Lucas, M. et al., 2008. A molecular framework for light and gibberellin control of cell elongation. Nature, 451(7177), pp.480–484.

Du, Z. et al., 2010. agriGO: a GO analysis toolkit for the agricultural community. Nucleic Acids Research, 38, pp.W64–W70.

Duek, P.D. & Fankhauser, C., 2005. bHLH class transcription factors take centre stage in phytochrome signalling. Trends in Plant Science, 10(2), pp.51–54.

Efroni, I. et al., 2013. Regulation of Leaf Maturation by Chromatin-Mediated Modulation of Cytokinin Responses. Developmental Cell, 24(4), pp.438–445.

Eklund, D.M. et al., 2010. The Arabidopsis thaliana STYLISH1 protein acts as a transcriptional activator regulating auxin biosynthesis. The Plant cell, 22(2), pp.349–363.

Farrona, S. et al., 2011. Brahma Is Required for Proper Expression of the Floral Repressor FLC in Arabidopsis M. Grebe, ed. PLoS ONE, 6(3), p.e17997.

Feng, S. et al., 2008. Coordinated regulation of Arabidopsis thaliana development by light and gibberellins. Nature, 451(7177), pp.475–479.

Frick, G. et al., 2003. An Arabidopsis porB porC double mutant lacking light-dependent NADPH:protochlorophyllide oxidoreductases B and C is highly chlorophyll-deficient and developmentally arrested. The Plant journal, 35(2), pp.141–153.

Friml, J. et al., 2003. Efflux-dependent auxin gradients establish the apical-basal axis of Arabidopsis. Nature, 426(6963), pp.147–153.

Friml, J., Benková, E., et al., 2002. AtPIN4 mediates sink-driven auxin gradients and root patterning in Arabidopsis. Cell, 108(5), pp.661–673.

Friml, J., Wiśniewska, J., et al., 2002. Lateral relocation of auxin efflux regulator PIN3 mediates tropism in Arabidopsis. Nature, 415(6873), pp.806–809.

Fuentes, S. et al., 2012. Fruit growth in Arabidopsis occurs via DELLA-dependent and DELLA-independent gibberellin responses. The Plant cell, 24(10), pp.3982–3996.

Gaillochet, C. et al., 2017. Control of plant cell fate transitions by transcriptional and hormonal signals. eLife, 6. e30135

Galvao, V.C. et al., 2012. Spatial control of flowering by DELLA proteins in Arabidopsis thaliana. Development, 139(21), pp.4072–4082.

Gietz, R.D. & Schiestl, R.H., 2007. High-efficiency yeast transformation using the LiAc/SS carrier DNA/PEG method. Nature Protocols, 2(1), pp.31–34.

Gremski, K., Ditta, G. & Yanofsky, M.F., 2007. The HECATE genes regulate female reproductive tract development in Arabidopsis thaliana. Development, 134(20), pp.3593–3601.

Groszmann, M. et al., 2011. SPATULA and ALCATRAZ, are partially redundant, functionally diverging bHLH genes required for Arabidopsis gynoecium and fruit development. The Plant journal, 68(5), pp.816–829.

Groszmann, M., Paicu, T. & Smyth, D.R., 2008. Functional domains of SPATULA, a bHLH transcription factor involved in carpel and fruit development in Arabidopsis. The Plant journal, 55(1), pp.40–52.

Hardtke, C.S. & Berleth, T., 1998. The Arabidopsis gene MONOPTEROS encodes a transcription factor mediating embryo axis formation and vascular development. The EMBO Journal, 17(5), pp.1405–1411.

Heisler, M.G. et al., 2001. SPATULA, a gene that controls development of carpel margin tissues in Arabidopsis, encodes a bHLH protein. Development, 128(7), pp.1089–1098.

Hoffman, C.S. & Winston, F., 1987. A ten-minute DNA preparation from yeast efficiently releases autonomous plasmids for transformation of Escherichia coli. Gene, 57(2-3), pp.267–272.

Hu, Z. et al., 2004. VisANT: an online visualization and analysis tool for biological interaction data. BMC Bioinformatics, 5, p.17.

Khanna, R. et al., 2004. A novel molecular recognition motif necessary for targeting photoactivated phytochrome signaling to specific basic helix-loop-helix transcription factors. The Plant cell, 16(11), pp.3033–3044.

Klepikova, A.V. et al., 2016. A high resolution map of the Arabidopsis thalianadevelopmental transcriptome based on RNA-seq profiling. The Plant Journal, 88(6), pp.1058–1070.

Kondoh, H. & Kamachi, Y., 2010. SOX-partner code for cell specification: Regulatory target selection and underlying molecular mechanisms. The international journal of biochemistry & cell biology, 42(3), pp.391–399.

Kubota, A. et al., 2017. TCP4-dependent induction of CONSTANS transcription requires GIGANTEA in photoperiodic flowering in Arabidopsis S. Hake, ed. PLoS Genetics, 13(6), p.e1006856.

Kuusk, S. et al., 2002. STY1 and STY2 promote the formation of apical tissues during Arabidopsis gynoecium development. Development, 129(20), pp.4707–4717.

Lampropoulos, A. et al., 2013. GreenGate---a novel, versatile, and efficient cloning system for plant transgenesis. PLoS ONE, 8(12), p.e83043.

Lau, O.S. et al., 2014. Direct roles of SPEECHLESS in the specification of stomatal self-renewing cells. Science, 345(6204), pp.1605–1609.

Leivar, P. & Quail, P.H., 2011. PIFs: pivotal components in a cellular signaling hub. Trends in Plant Science, 16(1), pp.19–28.

Liscum, E. & Briggs, W.R., 1995. Mutations in the NPH1 locus of Arabidopsis disrupt the perception of phototropic stimuli. The Plant cell, 7(4), pp.473–485.

Lucero, L.E. et al., 2015. TCP15 modulates cytokinin and auxin responses during gynoecium development in Arabidopsis. The Plant journal, 84(2), pp.267–282.

Martínez-Fernández, I. et al., 2014. The effect of NGATHA altered activity on auxin signaling pathways within the Arabidopsis gynoecium. Frontiers in plant science, 5, p.210.

Mathieu, J. et al., 2009. Repression of flowering by the miR172 target SMZ. PLoS Biology, 7(7), p.e1000148.

Mayer, K.F. et al., 1998. Role of WUSCHEL in regulating stem cell fate in the Arabidopsis shoot meristem. Cell, 95(6), pp.805–815.

Medzihradszky, A., Schneitz, K. & Lohmann, J.U., 2014. Detection of mRNA Expression Patterns by Nonradioactive In Situ Hybridization on Histological Sections of Floral Tissue. In J. L. Riechmann & F. Wellmer, eds. Flower Development: Methods and Protocols. Flower Development: Methods and Protocols. New York, NY: Springer New York, pp. 275–293.

Mering, von, C., 2004. STRING: known and predicted protein-protein associations, integrated and transferred across organisms. Nucleic Acids Research, 33, pp.D433–D437.

Moubayidin, L. & Østergaard, L., 2014. Dynamic Control of Auxin Distribution Imposes a Bilateral-to-Radial Symmetry Switch during Gynoecium Development. Current Biology, 24(22), pp.2743–2748.

Ni, M., Tepperman, J.M. & Quail, P.H., 1998. PIF3, a phytochrome-interacting factor necessary for normal photoinduced signal transduction, is a novel basic helix-loop-helix protein. Cell, 95(5), pp.657–667.

Paddock, T. et al., 2012. Arabidopsis light-dependent protochlorophyllide oxidoreductase A (PORA) is essential for normal plant growth and development. Plant Molecular Biology, 78(4-5), pp.447–460.

Peng, J. et al., 1997. The Arabidopsis GAI gene defines a signaling pathway that negatively regulates gibberellin responses. Genes & Development, 11(23), pp.3194–3205.

Pfeiffer, A. et al., 2014. Combinatorial complexity in a transcriptionally centered signaling hub in Arabidopsis. Molecular Plant, 7(11), pp.1598–1618.

Rajani, S. & Sundaresan, V., 2001. The Arabidopsis myc/bHLH gene ALCATRAZ enables cell separation in fruit dehiscence. Current Biology, 11(24), pp.1914–1922.

Reyes-Olalde, J.I. et al., 2017. The bHLH transcription factor SPATULA enables cytokinin signaling, and both activate auxin biosynthesis and transport genes at the medial domain of the gynoecium. PLoS Genetics, 13(4), p.e1006726.

Reymond, M.C. et al., 2012. A light-regulated genetic module was recruited to carpel development in Arabidopsis following a structural change to SPATULA. The Plant cell, 24(7), pp.2812–2825.

Roig-Villanova, I. et al., 2007. Interaction of shade avoidance and auxin responses: a role for two novel atypical bHLH proteins. The EMBO Journal, 26(22), pp.4756–4767.

Schaller, G.E., Bishopp, A. & Kieber, J.J., 2015. The yin-yang of hormones: cytokinin and auxin interactions in plant development. The Plant cell, 27(1), pp.44–63.

Schmid, M. et al., 2005. A gene expression map of Arabidopsis thaliana development. Nature Genetics, 37(5), pp.501–506.

Schoof, H. et al., 2000. The stem cell population of Arabidopsis shoot meristems in maintained by a regulatory loop between the CLAVATA and WUSCHEL genes. Cell, 100(6), pp.635–644.

Schuster, C. et al., 2014. A Regulatory Framework for Shoot Stem Cell Control Integrating Metabolic, Transcriptional, and Phytohormone Signals. Developmental Cell, 28(4), pp.438–449.

Schuster, C., Gaillochet, C. & Lohmann, J.U., 2015. Arabidopsis HECATE genes function in phytohormone control during gynoecium development. Development, 142(19), pp.3343–3350.

Serrano-Mislata, A. et al., 2017. DELLA genes restrict inflorescence meristem function independently of plant height. Nature Plants, 3(9), pp.749–754.

Silverstone, A.L. et al., 2001. Repressing a repressor: gibberellin-induced rapid reduction of the RGA protein in Arabidopsis. The Plant cell, 13(7), pp.1555–1566.

Silverstone, A.L., Ciampaglio, C.N. & Sun, T., 1998. The Arabidopsis RGA gene encodes a transcriptional regulator repressing the gibberellin signal transduction pathway. The Plant cell, 10(2), pp.155–169.

Smith, H., 2000. Phytochromes and light signal perception by plants--an emerging synthesis. Nature, 407(6804), pp.585–591.

Szklarczyk, D. et al., 2015. STRING v10: protein-protein interaction networks, integrated over the tree of life. Nucleic Acids Research, 43(D1), pp.D447–D452.

Tognetti, V.B. et al., 2010. Perturbation of indole-3-butyric acid homeostasis by the UDP-glucosyltransferase UGT74E2 modulates Arabidopsis architecture and water stress tolerance. The Plant cell, 22(8), pp.2660–2679.

Toledo-Ortiz, G., Huq, E. & Quail, P.H., 2003. The Arabidopsis basic/helix-loop-helix transcription factor family. The Plant cell, 15(8), pp.1749–1770.

Trigueros, M. et al., 2009. The NGATHA genes direct style development in the Arabidopsis gynoecium. The Plant cell, 21(5), pp.1394–1409.

Valverde, F. et al., 2004. Photoreceptor regulation of CONSTANS protein in photoperiodic flowering. Science, 303(5660), pp.1003–1006.

Vercruyssen, L. et al., 2014. ANGUSTIFOLIA3 binds to SWI/SNF chromatin remodeling complexes to regulate transcription during Arabidopsis leaf development. The Plant cell, 26(1), pp.210–229.

Weijers, D. & Wagner, D., 2016. Transcriptional Responses to the Auxin Hormone. Annual Review of Plant Biology, 67, pp.539–574.

Wigge, P.A. et al., 2005. Integration of spatial and temporal information during floral induction in Arabidopsis. Science, 309(5737), pp.1056–1059.

Wu, M.-F. et al., 2015. Auxin-regulated chromatin switch directs acquisition of flower primordium founder fate. eLife, 4, p.e09269.

Xu, X. et al., 2015. Illuminating Progress in Phytochrome-Mediated Light Signaling Pathways. Trends in Plant Science, 20(10), pp.641–650.

Yadav, R.K. et al., 2011. WUSCHEL protein movement mediates stem cell homeostasis in the Arabidopsis shoot apex. Genes & Development, 25(19), pp.2025–2030.

Zhou, P. et al., 2014. Both PHYTOCHROME RAPIDLY REGULATED1 (PAR1) and PAR2 Promote Seedling Photomorphogenesis in Multiple Light Signaling Pathways. Plant Physiology, 164(2), pp.841–852.

Zhu, L. et al., 2016. A Negative Feedback Loop between PHYTOCHROME INTERACTING FACTORs and HECATE Proteins Fine-Tunes Photomorphogenesis in Arabidopsis. The Plant cell, 28(4), pp.855–874.

Zürcher, E. et al., 2013. A robust and sensitive synthetic sensor to monitor the transcriptional output of the cytokinin signaling network in planta. Plant Physiology, 161(3), pp.1066–1075.

